# Intra-axonal translation of *Khsrp* mRNA slows axon regeneration by destabilizing localized mRNAs

**DOI:** 10.1101/2020.10.26.356162

**Authors:** Priyanka Patel, Courtney Buchanan, Amar N. Kar, Seung Joon Lee, Pabitra K. Sahoo, Anatoly Urisman, Juan Oses-Prieto, Michela Dell’Orco, Devon E. Cassidy, Sharmina Miller, Elizabeth Thames, Terika P. Smith, Matthew D. Zdradzinski, Alma L. Burlingame, Nora Perrone-Bizzozero, Jeffery L. Twiss

**Affiliations:** Department of Biological Sciences, University of South Carolina, Columbia, South Carolina 29208, USA; Department of Pharmaceutical Sciences, University of California, San Francisco, California 94143, USA; Department of Neurosciences, University of New Mexico School of Health Sciences, Albuquerque, New Mexico 87131, USA

**Keywords:** Axonal translation, mRNA stability, RNA binding protein, axon regeneration, conditioning nerve lesion

## Abstract

Proteins generated by localized mRNA translation in axons support nerve regeneration through retrograde injury signaling and localized axon growth mechanisms. RNA binding proteins (RBP) are needed for this and other aspects of post-transcriptional control of localized mRNAs, but only a limited number of axonal RBPs have been reported. We used a targeted mass spectrometry approach to profile the axonal RBPs in naïve, injured and regenerating PNS axons. We detected 76 axonal proteins that are reported to have RNA binding activity, with the levels of several of these axonal RBPs changing with axonal injury and regeneration. These axonal RBPs with altered axoplasm levels include KHSRP that we previously reported decreases neurite outgrowth in developing CNS neurons. We show that KHSRP levels rapidly increase in sciatic nerve axons after crush injury and remain elevated increasing in levels out to 28 days post-sciatic nerve crush injury. *Khsrp* mRNA localizes into axons and the rapid increase in axonal KHSRP after axotomy is mediated by the local translation of its mRNA. KHSRP binds to mRNAs with a 3’UTR AU-rich element and targets those mRNAs to the cytoplasmic exosome for degradation. KHSRP knockout mice show increased axonal levels of defined KHSRP target mRNAs, *Gap43* and *Snap25* mRNAs, following sciatic nerve injury and accelerated nerve regeneration *in vivo*. These data indicate that axonal translation of *Khsrp* mRNA following nerve injury serves to destabilize other axonal mRNAs and slow axon regeneration.

## INTRODUCTION

Localization of mRNAs to subcellular regions provides polarized cells with means to rapidly respond to environmental stimuli within different domains of the cell. Neurons are highly polarized cells with cytoplasmic processes, axons and dendrites, that extend great distances from the cell body or soma. Several lines of evidence indicate that translation products of mRNAs localizing into axons of the peripheral nervous system (PNS) contribute to axon regeneration after injury (Smith et al., 2020). In rodents, PNS and some CNS axons extend centimeters from their soma, so this localized protein synthesis provides autonomy from the soma but also must be tightly regulated. Since one mRNA can be translated many times over to generate multiple copies of a protein, the survival of an mRNA within an axon can substantially affect the spatial and temporal regulation of that axon’s proteome as well as its regeneration capacity (Dalla Costa et al., 2020). Much has been learned about how mRNAs are transported into and translated within axons over recent years, and it is clear that the axonal transcriptome is quite extensive in terms of numbers of different mRNAs and dynamic in terms of changes in mRNA populations with different physiological states (Kar et al., 2018). RNA binding proteins (RBP) and mRNAs assemble into a ribonucleoprotein particle (RNP) for transport into axons, and those or other RBPs can also subsequently regulate the translation of the mRNA in the axons or provide a storage depot to sequester the mRNA until needed (Dalla Costa et al., 2020). Both nonsense-mediated decay (NMD) and microRNA (miRNA)-stimulated RNA degradation have been shown to occur in distal axons (Aschrafi et al., 2012; Colak et al., 2013; Hengst et al., 2006; Murashov et al., 2007; Vargas et al., 2016). These observations point to an intrinsic capacity for locally depleting specific mRNAs from axons.

RBP interactions have also been linked to mRNA stability, including axonally localized mRNAs. The neuronal protein HuD (also called ELAVL4) has been known to stabilize mRNAs by binding to AU-rich elements (AREs) in 3’UTRs of target mRNAs (Beckel-Mitchener et al., 2002). HuD protein localizes into distal neurites and has been implicated in transport and translation of some mRNAs (Bolognani et al., 2004; Smith et al., 2004; Yoo et al., 2013). We previously showed that the KH splicing regulatory protein (KHSRP; also known as KSRP, MARTA1, ZBP2, and FUBP2) decreases neurite growth in cultures of embryonic cortical neurons and competes with HuD for binding to target mRNAs (Bird et al., 2013). In contrast to HuD interactions, KHSRP binding destabilizes ARE-containing mRNAs (Gherzi et al., 2004). Here, we show that KHSRP is one of several RBPs whose levels increase in PNS axons after injury and during regeneration. This increase in axonal KHSRP occurs rapidly after PNS nerve injury through translation of its mRNA in axons. Deletion of the KHSRP gene increases axonal levels of mRNAs encoding growth-associated proteins and this effect requires KHSRP’s fourth hnRNP K homology (KH) domain that was shown to bind to ARE-containing mRNAs (Gherzi et al., 2004). Our data indicate that KHSRP slows axon growth through axon-intrinsic mechanisms, and point to autonomy of the axonal compartment for modulating levels of axonal mRNAs and, thereby, the axon’s growth potential.

## RESULTS

### Peripheral nerve injury changes the axonal RNA binding protein population

Protein synthesis in PNS axons has been shown to facilitate nerve regeneration after injury. Transport and translation of the mRNA templates needed for this intra-axonal protein synthesis are driven by RBPs bound to those mRNAs (Dalla Costa et al., 2020). We recently showed that PNS axoplasm contains many RBPs that were thought to have exclusively nuclear roles by using RNA affinity mass spectrometry (RAMS) (Lee et al., 2018). The levels of some of these RBPs increased after axotomy, which pointed to functions in growing axons. As this RAMS assay focused on interactions with axonal mRNA localization motifs, we wanted to gain a more systematic and unbiased view of the axonal RBP population. With the limited amount of proteins obtained from PNS axoplasm coupled with the likely low abundance of RBPs relative to cytoskeletal components in the axons, we turned to a targeted mass spectrometry (MS) approach with parallel reaction monitoring (PRM) to profile axonal RBPs in naïve, injured and regenerating sciatic nerve axoplasm. By mining the *UniProt* database (https://www.uniprot.org) for proteins with ‘RNA binding’ in functional or domain descriptions and validated expression in the nervous system (cross-referencing each for RNA expression using the *GeneCards* database (https://www.genecards.org)), we arrived at 357 proteins to test. Of these, 196 were represented in a reference MS library that had been generated from rat nervous system tissues (including sciatic nerve, DRG, and spinal cord) and were included in a PRM-MS method that targeted 511 unique precursor peptides (Sahoo et al., 2018). Using this PRM method, we quantified the abundance of 84 RBPs represented by 184 precursor peptides in adult rat sciatic nerve axoplasm. Several of the proteins showed increases or decreases over 3-28 days post injury compared to the contralateral uninjured sciatic nerve (Figures 1A-B; Supplemental Figure S1). Initial validation by immunoblotting from naïve and 7 days post-crush injury sciatic nerve axoplasm showed that FXR1, hnRNP A3, hnRNP AB, hnRNP H1 and KHSRP increased in the 7 day samples (Figure 1C). Notably, hnRNP H1 was identified by the RAMS study for axonal RBPs binding to *Nrn1* mRNA localization motif and we similarly found its axoplasm levels are increased after injury (Lee et al., 2018).

**Figure 1:**
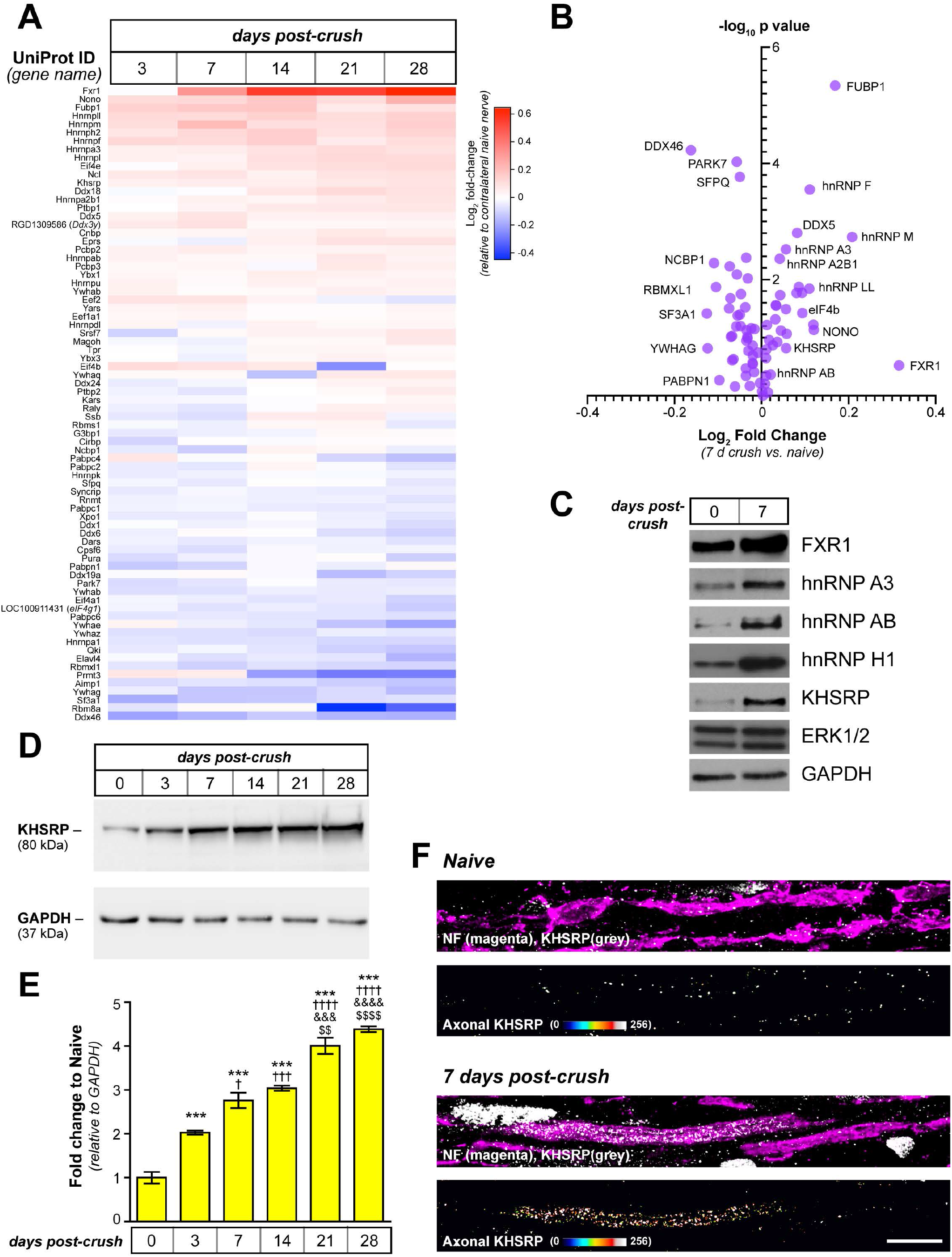
Peripheral nerve injury changes the axonal RNA binding protein populations. **A-B)** Sciatic nerve axoplasm harvested proximal to the injury site from 3-28 d post-crush lesion was digested and processed for liquid chromatography and mass spectrometry using parallel reaction monitoring (PRM) to detect proteins with known RNA binding activity. Levels of proteins from spectral counts relative to uninjured (naïve) axoplasm as shown as Log_2_ fold-change as indicated in **A** (N = 3 for each time point). **B** shows volcano plot for PRM results for 7 d crush vs. naïve samples graphed as Log_2_ fold-change vs. - Log_10_ p value for differences. A subset of proteins are identified here. **C)** Representative immunoblot for naïve and 7 d injured sciatic nerve axoplasm confirms the increase in FXR1, hnRNP A3, hnRNP AB, hnRNP H1, and KHSRP. ERK 1/2 and GAPDH show relatively equivalent loading of the lysates. Also see Suppl. Figure S1 for graphical representation of full time course. **D-E)** Representative immunoblots for kinetics of KHSRP elevation in sciatic nerve axoplasm over 0-28 d post-crush lesion are shown in **D**. Quantification of KHSRP immunoreactivity across multiple animals is shown as mean fold-change relative to naïve ± standard error of the mean (SEM; N = 3 for each time point; *** = P ≤ 0.001 for vs. 0 d, † = p ≤ 0.05, ††† = p ≤ 0.005, ††† = p ≤ 0.001 and †††† = p ≤ 0.0005 vs. 3 d, &&& = p ≤ 0.001 and &&&& = p ≤ 0.0005 vs. 7 d, and $$ = p ≤ 0.01 and $$$$ = p ≤ 0.0005 vs. 14 d by one way ANOVA with Tukey’s post-hoc analysis). **F)** Representative confocal images for KHSRP protein in naïve and 7 d post-crush sciatic nerve. Upper panels of each pair show XYZ maximum projection of merged KHSRP (grey) and neurofilament (NF; magenta) signals; lower panels show KHSRP signals overlapping with NF (‘axonal KHSRP’) in each individual Z plane projected as an XYZ image [Scale bar = 5 µm].

The increase in KHSRP after injury was surprising as the protein has been linked to mRNA destabilization in neurons, including targeting *Gap43* mRNA for degradation whose encoded protein has long been linked to axon growth promotion (Bird et al., 2013). Examining KHSRP more closely, its levels increased in the sciatic nerve axoplasm by 3 days (Fig 1D-E). Moreover, the axoplasm KHSRP levels were significantly higher at 21-28 days post injury than earlier time points (Fig 1D-E). With the mid-thigh sciatic nerve crush used, rats here begin to regain lower hind limb function over 21-28 days post injury indicative of some target reinnervation. The axoplasm preparation used is highly enriched for axonal proteins, but does contain some glial constituents (Rishal et al., 2010). Since KHSRP is ubiquitously expressed (Li et al., 2012; Min et al., 1997), we used confocal microscopy to determine if the elevations in KHSRP levels seen by PRM and immunoblotting derived from axonal KHSRP. Consistent with the ubiquitous KHSRP expression, KHSRP protein immunofluorescence was seen in both axons and the adjacent Schwann cells; however, extraction of the KHSRP immunofluorescence overlapping with neurofilament signals, showed a markedly increased axonal KHSRP signals at 7 days post injury compared to uninjured nerve (Figure 1F). Thus, axotomy induces an early and lasting increase in axonal KHSRP levels.

### Loss of KHSRP enhances axon growth in the peripheral nervous system through an axon-intrinsic mechanism

With its role in targeting mRNAs for degradation and clear evidence indicating that axonal mRNA translation supports axon regeneration, we contemplated what function this increased axonal KHSRP might serve. To address this, we took advantage of a global KHSRP knockout mouse line, where both alleles of KHSRP are deleted (Li et al., 2012). Cultures of dissociated lumbar (L) 4-6 dorsal root ganglia (DRGs) neurons from *Khsrp*^*-/-*^ mice showed significantly increased axon growth compared to those from *Khsrp*^*+/+*^ mice (Figure 2A-B) but no differences in axon branching (Suppl. Figure S2A-B).

**Figure 2:**
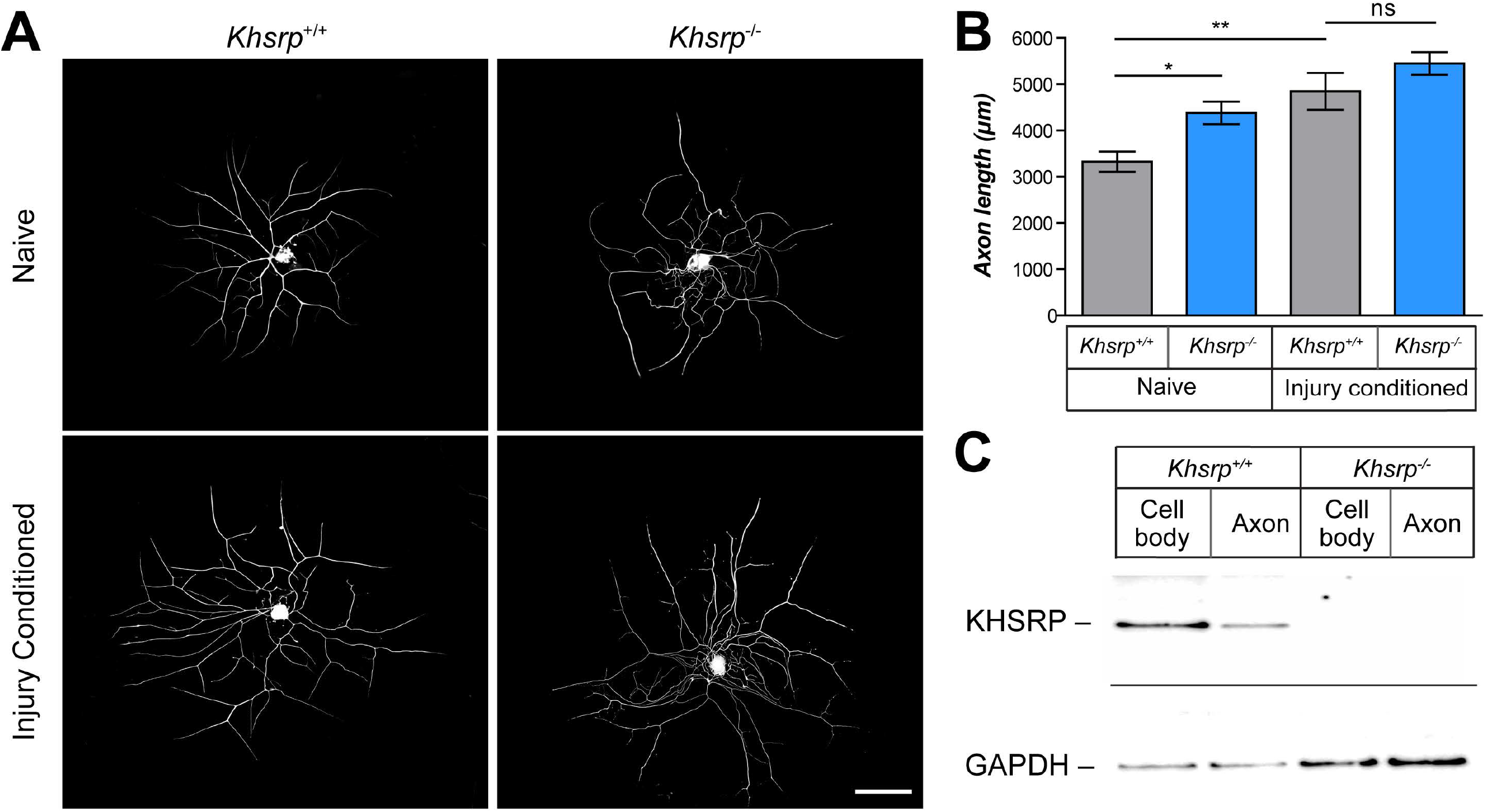
KHSRP depletion enhances axonal growth from naïve but not injury-conditioned DRG neurons. **A)** Representative images of 24 h DRG neuron cultures from naïve and 7 d injury-conditioned *Khsrp*^*+/+*^ and *Khsrp*^*-/-*^ mice immunostained for NF are shown [Scale bar = 100 µm]. **B)** Quantification of total axon length per neuron from cultures as in A shows significantly greater axon growth in *Khsrp*^*-/-*^ compared to *Khsrp*^*+/+*^ DRGs. However, there is no significant difference in axon length between the genotypes when cultures were prepared 7 d after nerve crush injury (*i*.*e*., ‘injury-conditioned’ neurons). Data are expressed as mean ± SEM; refer to Supplemental Figure S2 for axon branching analyses (≥ 75 neurons analyzed per sample over 3 independent cultures; * = P ≤ 0.05 and ** = P ≤ 0.01 by one way ANOVA with Tukey’s post-hoc analysis). **C)** Representative immunoblot shows signal for KHSRP in cell body and axonal preparations from DRG cultures from indicated mice. There is no KHSRP band in the *Khsrp*^*-/-*^ cell body or axonal preparations, despite GAPDH showing more protein in those samples than in the *Khsrp*^*+/+*^ DRGs.

*In vivo* injury-conditioning by sciatic nerve crush injury has been shown to increase *in vitro* axon growth from dissociated DRG neurons independent of new gene transcription upon culturing (Smith and Skene, 1997). The *in vivo* conditioning injury activates transcription of growth-associated genes, whose mRNA products are then translationally regulated after the second injury that is brought by the DRG dissection and dissociation needed for culture (Twiss et al., 2000b). Thus, we tested whether axon growth from ‘injury-conditioned’ L4-6 DRG s is further increased in the absence of KHSRP. In contrast to the naïve DRG cultures, there was no significant difference in axon length comparing the injury-conditioned DRGs from *Khsrp*^*-/-*^ vs. *Khsrp*^*+/+*^ mice (Figure 2A-B). To be certain that KHSRP was indeed absent from these neurons, we tested KHSRP levels in soma and axon lysates from L4-6 DRG neurons cultured neurons on a porous membrane for separation of axons (Willis and Twiss, 2011). KHSRP immunoreactive bands were clearly present in the *Khsrp*^*+/+*^ somal and axonal lysates, but were not detected in the *Khsrp*^*-/-*^ samples even with extended exposures (Figure 2C). These data suggest that removing KHSRP can increase axon growth, but maximum growth promotion compared to *Khsrp*^*+/+*^ DRG cultures was only seen in naïve and not injury-conditioned neurons.

With no differences in axon growth between in *Khsrp*^*-/-*^ and *Khsrp*^*+/+*^ injury-conditioned DRG neurons, this could suggest that the growth attenuating effects of KHSRP are overcome by the conditioning effects. However, since the MS data pointed to an increase of KHSRP within the axons of peripheral nerve after injury, an alternative hypothesis is that the growth-attenuating functions of KHSRP are axon-intrinsic. That is, shearing the DRG axons during *in vitro* culture preparation could negate any *in vivo* increase in KSHRP levels that is selective to the axons. To test this alternate hypothesis, we compared *in vivo* axon regeneration in the *Khsrp*^*-/-*^ and *Khsrp*^*+/+*^ mice. There was a modest increase in axon regeneration in the sciatic nerve regeneration at 7 days after crush injury in *Khsrp*^*-/-*^ vs. *Khsrp*^*+/+*^ mice, but this did not reach significance (Suppl. Figure S3). We next asked whether an *in vivo* injury-conditioning paradigm might fully distinguish the *Khsrp*^*-/-*^ and *Khsrp*^*+/+*^ mice, since the robust axotomy induced increase in axonal KHSRP would be absent in the *Khsrp*^*-/-*^ mice. Thus, we performed a unilateral sciatic nerve crush injury and 7 d later performed a second ipsilateral crush proximal to the initial injury; the contralateral side underwent single crush at the same time as the second lesion (and same approximate level). 3 d later, nerve regeneration contralateral to the conditioning lesion (*i*.*e*., single crush injury) showed no significant difference between *Khsrp*^*-/-*^ and *Khsrp*^*+/+*^ mice (Figure 3A-B). In sharp contrast, the injury-conditioned *Khsrp*^*-/-*^ nerves (*i*.*e*., double crush injury) showed a dramatic increase in axon regeneration compared to the injury-conditioned *Khsrp*^*+/+*^ mice (Figure 3A-B). To determine if this increased axon growth accelerated target reinnervation, we analyzed neuromuscular junctions (NMJs) in the gastrocnemius muscle 14 days after crush injury in *Khsrp*^*+/+*^ and *Khsrp*^*-/-*^ mice (Figure 3C). NMJ occupancy based on percentage of presynaptic compared postsynaptic area showed no differences between *Khsrp*^*+/+*^ and *Khsrp*^*-/-*^ mice at 14 days after a single nerve crush (Figure 3D). However, the double crush injured mice showed significantly greater NMJ occupancy in *Khsrp*^*-/-*^ vs. *Khsrp*^*+/+*^ (*i*.*e*., 14 d after the second crush lesion, 21 d after initial conditioning injury; Figure 3C-D).

**Figure 3:**
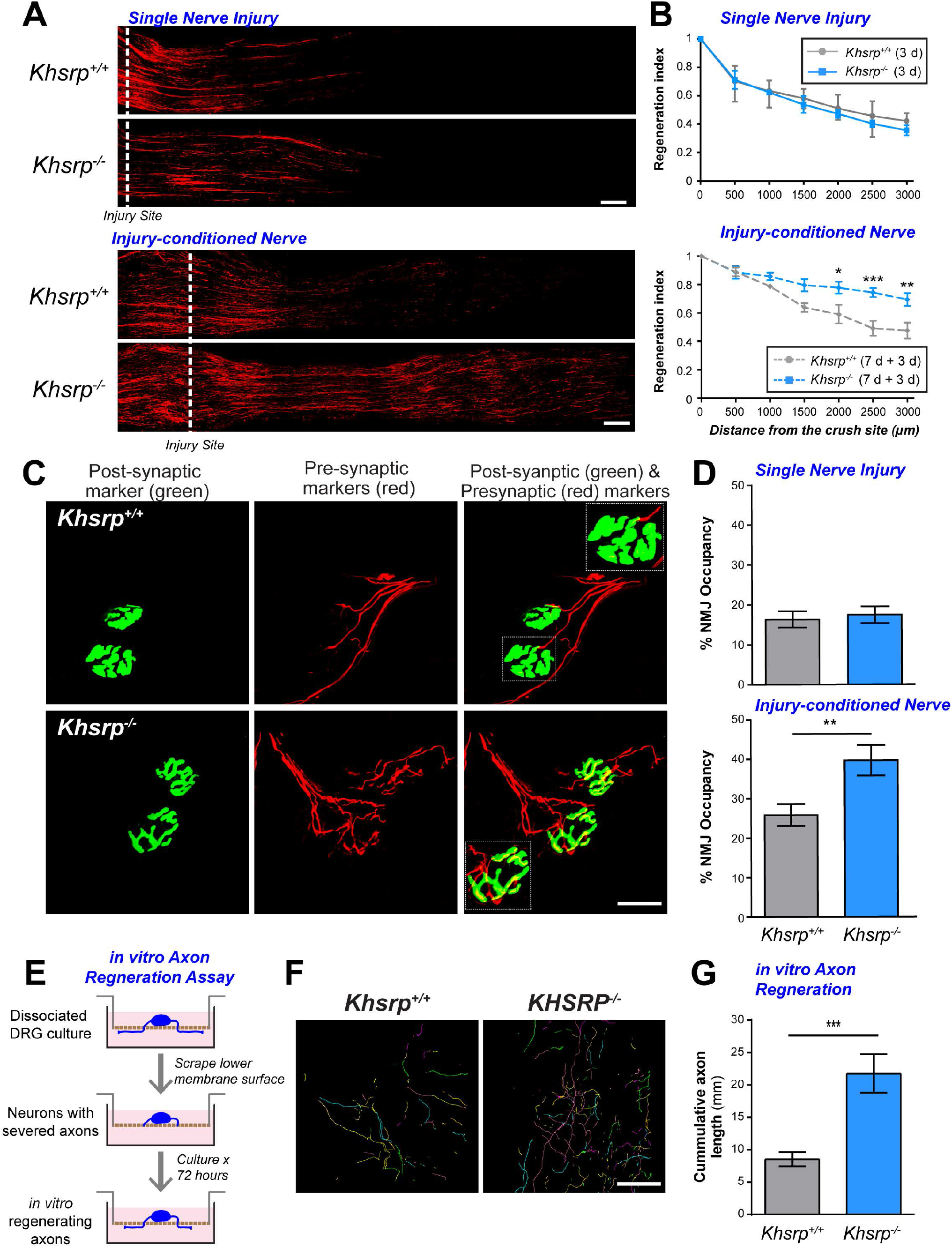
KHSRP deletion increases in vivo axon regeneration after a conditioning Sciatic nerve injury. **A)** Representative SCG10 immunostained images of sciatic nerves after single (3 d; upper panels) or injury-conditioned (7 + 3 d; lower panels) sciatic nerve crush injuries for *Khsrp*^*+/+*^ and *Khsrp*^*-/-*^ are shown. Proximal is on the left and distal on the right; the dashed line indicates the injury site, with the second injury for the double crush injured animals placed at ∼0.5 cm proximal to the initial injury site [Scale bar = 500 µm]. **B)** Regeneration indices calculated as fraction of SCG10 axonal profiles at injury site (0 µm) are shown as mean ± SEM as indicated. There was no significant difference in the regeneration after the single injury, but the injury-conditioned *Khsrp*^-/-^ mice show significantly higher regeneration indices (N = 5 mice per genotype and condition; * = P ≤ 0.05, ** = P ≤ 0.01, and *** = P ≤ 0.001 by TWO way ANOVA with Bonferroni post-hoc analysis). **C)** Confocal XYZ images of gastrocnemius muscles of the injury-conditioned *Khsrp*^*+/+*^ and *Khsrp*^*-/-*^ mice at 14 d after second nerve crush are shown. NMJs are detected by post-synaptic (*α*-bungarotoxin; green) and pre-synaptic markers (cocktail of anti-NF, -synapsin I, and -synaptophysin; red) signals showing higher matching of pre- and post-synaptic markers in *Khsrp*^*-/-*^ than *Khsrp*^*+/+*^ mice [Scale bar = 20 µm]. **D)** Quantification of NMJ occupancy (% presynaptic area/postsynaptic area) shows significantly greater occupancy in the injury-conditioned *Khsrp*^*-/-*^ mice but no difference with the single crush lesion (injury-conditioned = 7 + 14 d; single nerve crush = 14 d). Data are expressed as mean ± SEM (N ≥ 15 NMJs quantified in 3 animals per condition per genotype; ** = P ≤ 0.01 by Student’s t test). **E)** Schematic for the culture system to allow for axotomy and regeneration from the injured axon rather than initiation of axon growth from the cell body as seen in cultures of *in vivo* injury-conditioned neurons. For this, dissociated DRGs are grown on a porous membrane for 36 h. Axons are then severed by scraping the undersurface of the membrane. Axon regrowth along the undersurface of the membrane was assessed 72 h later. **F)** Representative *Neurolucida* tracings for NF-stained axons from filter undersurface for *Khsrp*^*+/+*^ and *Khsrp*^*-/-*^ DRGs at 72 h post *in vitro* axotomy [Scale bar = 100 µm]. **G)** Quantification of summed axon lengths per filter from G are shown as mean ± SEM. The *Khsrp*^*-/-*^ DRG cultures show significantly more *in vitro* axon regeneration than *KHSRP*^*+/+*^ DRG cultures (N = 3 for each genotype with ≥ 17 random fields analyzed; *** = P ≤ 0.001 by Student’s *t-*test).

The differences seen above between *in vivo* nerve regeneration and *in vitro* axon growth could be consistent with axon-intrinsic effects of KHSRP. To test this possibility more directly, we developed an *in vitro* axotomy model where DRGs were initially allowed to extend axons and then the axon shafts were severed. This allowed us to assess regeneration *in vitro* from an injured axon rather than initiation of axon growth from the cell body reflected in the *in vivo* injury-conditioned DRG cultures used in Figure 2B above or replated DRG neurons used in other studies (Perry et al., 2016). DRG neurons from *Khsrp*^*-/-*^ and *Khsrp*^*+/+*^ mice were cultured on membrane filters for 36 h; the undersurface of the membrane was then scraped to sever the axons (Figure 3E). After an additional 72 h in culture, neurofilament immunostained axons were traced along the membrane undersurface and total axon lengths were quantified (Figure 3F). Consistent with increased regeneration from injury-conditioned nerves *in vivo*, the *Khsrp*^*-/-*^ DRGs showed significantly accelerated axon regeneration after their axon shafts were severed *in vitro* (Figure 3G). Taken together, these data indicate that the intra-axonal increase in KHSRP following axotomy actually slows growth of the regenerating axon.

### The axotomy-induced increase in axonal KHSRP occurs via axon-intrinsic translation

Translation of mRNA in axons provides a means to rapidly change protein content in subcellular domains. For example, injury-induced translation of axonal *Calr* mRNA was recently shown to support early regrowth of severed axons (Pacheco et al., 2020). Thus, we wondered whether the change in axonal KHSRP levels might be driven by localized translation of its mRNA. We had previously detected *Khsrp* mRNA in RNA-seq analyses of sciatic nerve axoplasm (Lee et al., 2018); however, these axoplasm preparations obviously can contain non-neuronal contents (Rishal et al., 2010), particularly for mRNAs isolated from injured nerve as Lee *et al*. (2018) used. To overcome this limitation, we used single molecule fluorescence *in situ* hybridization (smFISH) combined with IF to ask whether sensory axons contain *Khsrp* mRNA. Dissociated DRG cultures from *Khsrp*^*+/+*^ mice showed prominent *Khsrp* mRNA signals in cell body and axons (Figure 4A-C). The axonal RNA signal was granular (Figure 4A), as typically seen for axonal and dendritic mRNAs. There was no *Khsrp* mRNA signal in the axons of the *Khsrp*^*-/-*^ DRG cultures (Figure 4C). Similarly, smFISH/IF performed on sciatic nerve sections showed prominent *Khsrp* mRNA signal in the axons for the *Khsrp*^*+/+*^ mice that was completely absent in *Khsrp*^*-/-*^ mice (Figure 4D-E). FISH signals for the cell bodies of the DRGs from *Khsrp*^*-/-*^ mice did show a faint but consistent signal with the *Khsrp* mRNA probes in the cell bodies that was significantly greater than scrambled probe (Figure 4A-B). The *Khsrp*^*-/-*^ mice were generated by deleting exons 1-13 (of 18) of the murine *KHSRP* gene (Lin et al., 2011), and the *Stellaris* smFISH *Khsrp* mRNA probes used here hybridize to sequences across the full exons comprising the mature *Khsrp* mRNA. Thus, with absence of KHSRP protein in these cultures based on Figure 2C, we suspect that this is an RNA transcribed from exons 14-18 remaining in the *Khsrp* gene, or portions thereof as confirmed by RNA-sequencing of *Khsrp*^*-/-*^ adult mouse brain and embryonic cortical neuron cultures (data not shown). Nonetheless, the strong *Khsrp* mRNA signal in the axons of cultured DRG neurons and *in vivo* sciatic nerve indicate that *Khsrp* mRNA localizes into PNS axons.

**Figure 4:**
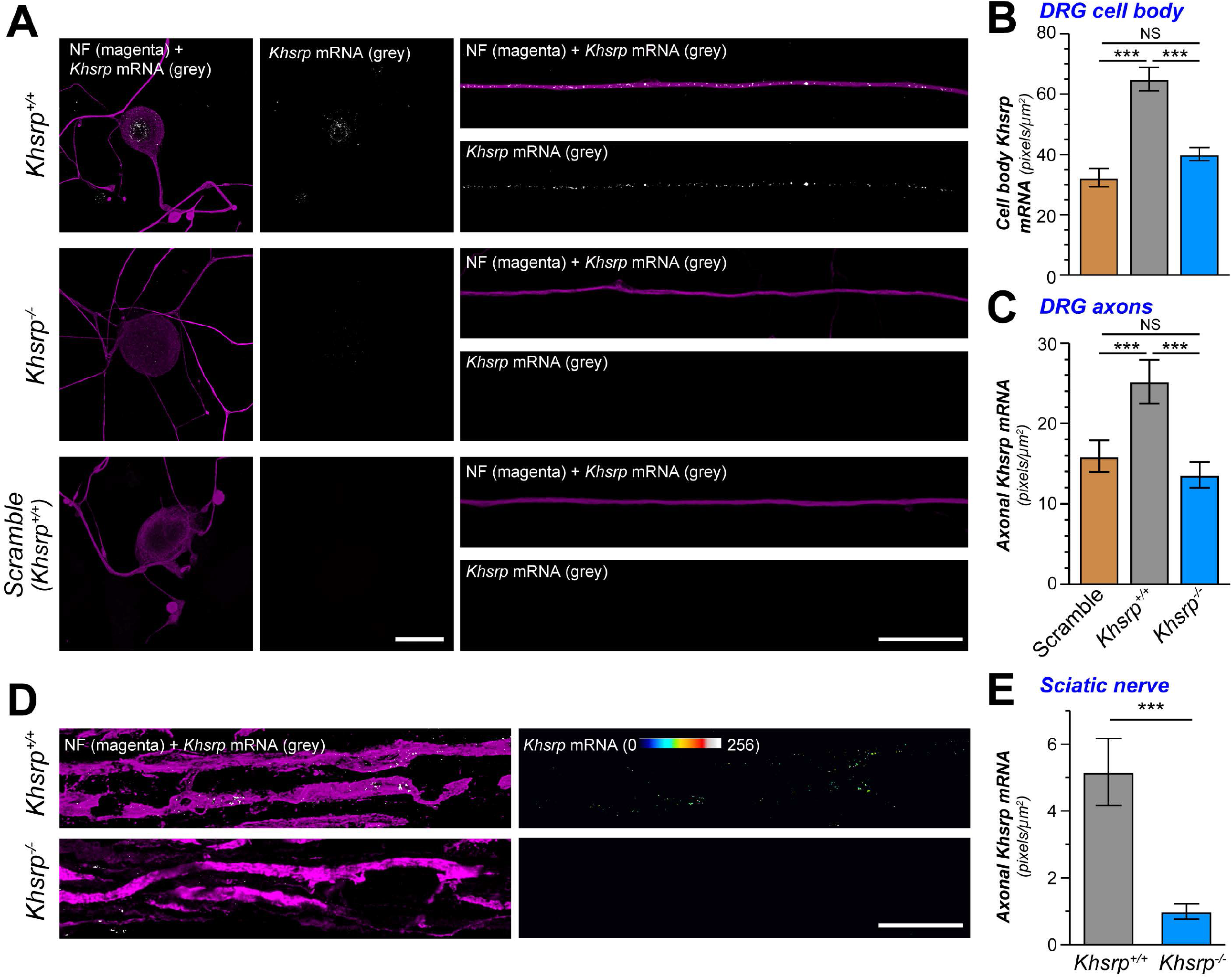
Khsrp mRNA is transported into PNS axons. **A)** Representative confocal images for smFISH/IF for *Khsrp* mRNA (grey) and NF (magenta) in dissociated DRG cultures are shown as indicated. There is a clear signal for *Khsrp* mRNA in the cell body (left and middle columns) and distal axons (right column) of DRGs from *Khsrp*^*+/+*^ mice. *Khsrp*^*-/-*^ DRGs showed no *Khsrp* mRNA in the axons; however, there was faint but consistent signal in the cell body [Scale bar = 10 µm]. **B-C)** Quantification smFISH signals for *Khsrp* mRNA in cell bodies (**B**) and axons (**C**) is shown as mean ± SEM for scramble (*Khsrp*^*+/+*^) and *Khsrp* mRNA (*Khsrp*^*+/+*^ and *Khsrp*^*-/-*^) probes; in each case, scramble probe was hybridized to *Khsrp*^*+/+*^ DRG cultures as in panel A (N ≥ 16 neurons over three separate cultures; * = P ≤ 0.05, ** = P ≤ 0.01, and *** = P ≤ 0.001 by one way ANOVA with Tukey’s post-hoc analysis). **D)** Representative confocal images for smFISH/IF for *Khsrp* mRNA (grey) and NF (magenta) in uninjured sciatic nerve are shown as indicated. Left column shows XYZ projections from eight optical planes taken at 0.2 µm Z step interval; right column shows ‘axon only’ *Khsrp* mRNA signals generated by extracting FISH signals overlapping with NF in individual Z sections and projecting those as an ‘Axonal *Khsrp* mRNA’ XYZ image [Scale bar = 5 µm]. **E)** Quantification of axonal *Khsrp* mRNA signals from D are shown as mean ±SEM (N = 6 animals per genotype; ** = P ≤ 0.01 by Student’s *t*-test).

The RNA analyses above suggest that KHSRP may be synthesized locally in axons. To determine if the PNS nerve injury-induced increase in KHSRP is intrinsic to the nerve rather than transported from cell body, we ligated the sciatic nerve to restrict anterograde transport and performed a crush injury distal to the ligation (Figure 5A-B). Adult rats were used for these ligation analyses since the larger sciatic nerve could be ligated more consistently than in the mouse. Immunofluorescence for amyloid precursor protein (APP) and Signal transducer and activator of transcription 3*α* (Stat3*α*) confirmed that the ligation attenuated both anterograde and retrograde transport, as APP accumulated proximal to and Stat3*α* accumulated distal to the ligation site (Suppl. Figure S4A). As expected KHSRP signals were detected both in axons and adjacent Schwann cells (Figure 5B-C, Suppl. Figure 4B-C). Axonal KHSRP Immunofluorescence showed increased signals at both the ligation (proximal and distal) as well as at the crush site relative to axons in naïve nerve; however, the increased axonal KHSRP signals proximal to the crush site were significantly greater and continued to elevate over time after injury compared to the ligation sites (Figure 5B-C, Suppl. Figure 4B-C). These results are consistent with local synthesis of KHSRP in the injured nerve.

**Figure 5:**
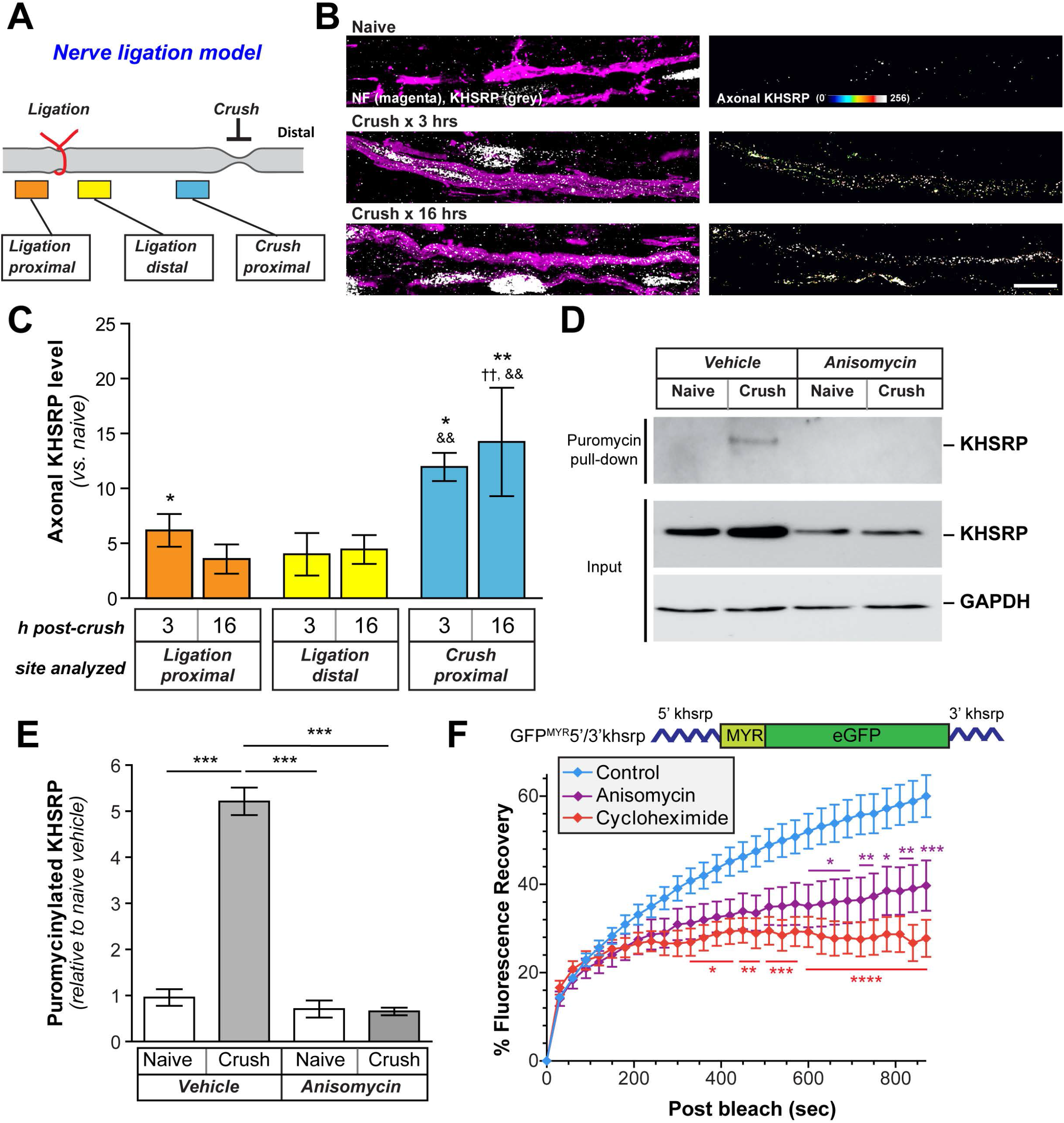
Khsrp mRNA is rapidly translated in axons after injury. **A)** Schematic of nerve ligation model used for panels B-C. Proximal nerve is on the left and distal on the right as indicated. The nerve was ligated and then immediately crushed ∼1 cm distal to the ligation. **B)** Confocal images for KHSRP protein in naïve and post-crush injury (3 and 16 h). Left column shows XYZ projections of merged signals for KHSRP (grey) and NF (magenta); right column shows ‘axonal KHSRP’ signals as from individual Z planes that were projected as an XYZ image. Representative images for ligation efficiency and KHSRP signals proximal and distal to ligation are shown in Suppl. Figure S4 [Scale bar = 5 µm]. **C)** Quantification of the axonal KHSRP immunoreactivity from ligation proximal and distal and crush sites are shown as mean ± SEM (N ≥ 3 animals per time point; * = p ≤ 0.05 and ** = p ≤ 0.01 for indicated time points vs. naïve nerve, †† = p ≤ 0.01 for indicated sample vs. crush proximal, and && = p ≤ 0.01 for indicated sample vs. crush distal by Student’s *t*-test; ligation proximal vs. distal have no significance). **D)** Representative immunoblots for *ex vivo* puromycinylated naïve vs. crushed sciatic nerve segments are shown as indicated. Protein synthesis inhibition with anisomycin completely blocks the puromycinylation of KHSRP in axoplasm samples extruded from the nerve segments, and GAPDH shows relatively equivalent protein loading across the lanes. **E)** Quantification of puromycinylated KHSRP signals from D is shown as mean ± SEM. Crush injury significantly increases axonal KHSRP synthesis and this blocked by anisomycin (N = 3; *** = P ≤ 0.001 by one way ANOVA with Tukey’s post-hoc analysis). **F)** FRAP analyses for distal axons of neurons transfected with GFP^MYR^5’/3’khsrp (see schematic at top) is shown as normalized average % recovery ± SEM. Pre-treatment treated with anisomycin or cycloheximide significantly attenuates the GFP recovery, indicating protein synthesis dependent recovery in the axons (N ≥ 30 neurons over 3 repetitions, * p ≤ 0.05, ** p ≤ 0.01, and *** p ≤ 0.005 by two-way ANOVA with Bonfieri post-hoc analyses for indicated time points vs. control). Representative images sequences for FRAP are shown in Suppl. Figure S5.

To more directly test if KHSRP is translated in the sciatic nerve axons, we exploited an *ex vivo* nerve injury model where we used puromycin incorporation to detect nascent peptides from the axoplasm extruded from the nerve segments (Terenzio et al., 2018). We excised segments of rat sciatic nerve and placed these into culture medium; cyclosporin A was included to delay Wallerian degeneration (Barrientos et al., 2011) and O-parpargyl-puromycin (OPP) was included for puromycinylation of nascently synthesized polypeptides (Forester et al., 2018; Sahoo et al., 2020). This *ex vivo* assay showed a significant increase in puromycinylated KHSRP in the axoplasm isolates at 4 h after nerve crush that was attenuated by the protein synthesis inhibitor anisomycin (Figure 5D-E). Notably, an increase in overall KHSRP levels was seen in the crushed nerve segment axoplasm and this was also attenuated by pretreatment with anisomycin (Figure 5D), confirming that axonal injury increases axonal KHSRP levels through mRNA translation. To further test for translation of *Khsrp* mRNA in axons, we fused 5’ and 3’UTRs of rodent *Khsrp* mRNA to the coding sequence of a diffusion-limited GFP reporter cDNA (GFP^MYR^5’/3’khsrp; Figure 5F) to use as a surrogate for axonal localization and translation of *Khsrp* mRNA. By fluorescence recovery after photobleaching (FRAP), DRG neurons expressing GFP^MYR^5’/3’khsrp showed rapid fluorescent recovery that was significantly attenuated by pretreatment with anisomycin or a second protein synthesis inhibitor cycloheximide (Figure 5F; Suppl. Figure S5). Taken together these results indicate that the axotomy-induced increase in axonal KHRP is derived from localized translation of *Khsrp* mRNA in axons.

### KHSRP’s ARE-binding KH4 domain attenuates axon growth

KHSRP was previously shown to bind to the ARE in *Gap43* mRNA’s 3’UTR and destabilize the transcript (Bird et al., 2013). The *GAP43* gene is transcriptionally activated following PNS nerve injury (Van der Zee et al., 1989), and axonal *Gap43* mRNA levels increase more than 2 fold in regenerating PNS axons (Yoo et al., 2013). Given that *Gap43* mRNA and protein expression are typically associated with axon growth (Skene, 1989), we tested the effect of *Khsrp* knockout on the sciatic nerve levels of this KHSRP target mRNA. RTddPCR from sciatic nerve samples of *Khsrp*^*+/+*^ mice showed the expected increase in *Gap43* mRNA at 7 days after nerve crush injury (Figure 6A). However, the sciatic nerve levels of *Gap43* mRNA in *Khsrp*^*-/-*^ mice were approximately 2-fold higher than in *Khsrp*^*+/+*^ mice after crush injury (Figure 6A). *Gap43* mRNA levels in injured *Khsrp*^*+/-*^ mouse nerves were intermediate between *Khsrp*^*+/+*^ and *Khsrp*^*-/-*^ nerves, but this did not reach statistical significance. There were only modest, non-significant increases in nerve *Gap43* mRNA for uninjured *Khsrp*^*-/-*^ and *Khsrp*^*+/-*^ vs. *Khsrp*^*+/+*^ nerves (Figure 6A). Since KHSRP can potentially bind to many different axonal mRNAs, we asked if axonal *Snap25* mRNA might be affected by *Khsrp* knockout. We recently found that *Snap25* mRNA is a target for destabilization by KHSRP in mouse brain(Olguin et al., 2020), and the Hengst lab had shown that translation of *Snap25* mRNA in CNS axons promotes synaptogenesis (Batista et al., 2017). *Snap25* mRNA was detected in sciatic nerve, and its levels were significantly increased in the injured sciatic nerves of *Khsrp*^*-/-*^ compared to *Khsrp*^*+/+*^ mice (Figure 6B). In contrast, there was no difference in the levels of *Hmgb1* mRNA (also called Amphoterin; Figure 6C), an mRNA that we have previously shown localizes to axons (Merianda et al., 2015) but was not affected by *Khsrp* knockout nor did it bind to KHSRP in RIP-Seq analyses (Olguin et al., 2020). Thus, increased levels of mRNAs encoding proteins linked to axon growth accompany the increased axon regeneration seen in injury-conditioned *Khsrp*^-/-^ mice.

**Figure 6:**
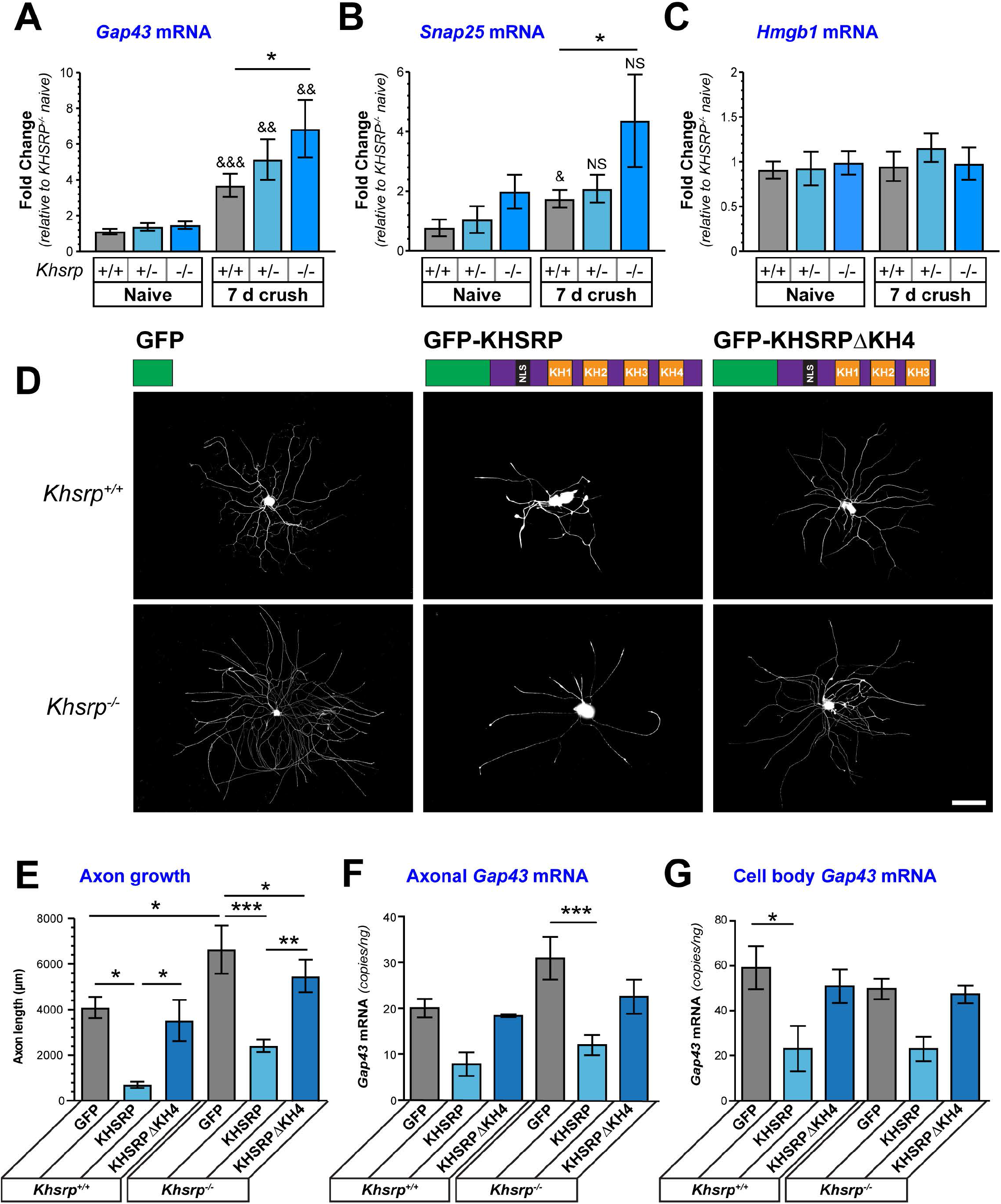
The ARE-binding KH4 domain of KHSRP attenuates axon growth and axonal mRNA levels. **A-C)** Sciatic nerve levels of *Gap43* (**A**) and *Snap25* (**B**) mRNA levels are increased 7 d after sciatic nerve crush in *Khsrp*^-/-^ compared to *Khsrp*^*+/+*^ mice. Sciatic nerve *Hmgb1* mRNA levels (**C**) show no change comparing the *Khsrp*^-/-^ vs. *Khsrp*^*+/+*^ mice or with injury. All genotypes show a significant increase for *Gap43* mRNA with crush injury. Values shown as mean ± SEM (N = 5 mice per genotype; * = P ≤ 0.05 for indicated conditions by two-way ANOVA with Bonferroni post-hoc; & = P ≤ 0.05, && = P ≤ 0.01, and &&& = P ≤ 0.005 and NS = not significant for crush vs. naïve within genotype by Student’s t-test). **D)** Representative images of DRG neurons transfected with GFP, GFP-KHSRP, or GFP-KHSRPΔKH4 are shown for *Khsrp*^*+/+*^ and *Khsrp*^*-/-*^ (schematics for constructs above image columns) as indicated [Scale bar = 100 µm]. **E)** Quantification of total axon length/neuron for DRG cultures transfected as in D are shown as mean ± SEM. Expression of GFP-KHSRP decreases axon outgrowth in both *Khsrp*^*+/+*^ and *Khsrp*^*-/-*^ DRGs, but GFP-KHSRPΔKH4 had no significant effect on *Khsrp*^*+/+*^ and only a modest decrease in axon length in *Khsrp*^*-/-*^ neurons (N ≥ 30 neurons over three different culture preparations; * = P ≤ 0.05, ** = P ≤ 0.01, and *** = P ≤ 0.001 by one-way ANOVA with Tukey’s post-hoc analysis for indicated comparisons). **F-G)** Analysis of axonal (**F**) and cell body (**G**) *Gap43* mRNA levels in neurons transfected as in D-E is shown as mean mRNA copies/ng of total RNA ± SEM after normalization to mitochondrial 12S RNA (N = 3 per genotype and per condition; * = P ≤ 0.05 and *** = P ≤ 0.001 by one-way ANOVA with Tukey’s post-hoc analysis for indicated comparisons).

Previous work has shown that the KH3 and KH4 domains of KHSRP are needed for ARE binding and mRNA degradation via the cytoplasmic exosome complex (Gherzi et al., 2004), and deletion of KH4 attenuates the effects of KHSRP on neurite growth in embryonic CNS neuron cultures (Bird et al., 2013). To determine if KHSRP regions previously linked to mRNA decay determine the axonal growth-attenuation by KHSRP in adult neurons, we transfected *Khsrp*^*+/+*^ and *Khsrp*^*-/-*^ DRG cultures with full length GFP-KHSRP or GFP-KHSRP with KH4 domain deleted (GFP-KHSRPΔKH4). GFP-KHSRP reversed the axon growth promoting effects of KHSRP loss in the *Khsrp*^-/-^ DRGs and decreased axon growth in *Khsrp*^*+/+*^ DRGs (Figure 6D-E). In contrast, GFP-KHSRP*Δ*KH4 expression only modestly decreased axon growth in the *Khsrp*^*-/-*^ DRGs and had no significant effect on *Khsrp*^*+/+*^ DRGs (Figure 6D-E). Axonal and cell body *Gap43* mRNA levels showed comparable results, with GFP-KHSRP expression significantly decreasing *Gap43* mRNA levels in *Khsrp*^*+/+*^ and *Khsrp*^*-/-*^ DRGs, while only modest or no decreases were seen with GFP-KHSRP*Δ*KH4 expression in both the *Khsrp*^*-/-*^ and *Khsrp*^*+/+*^ DRGs (Figure 6F-G). Taken together these data indicate that the RNA degradation activity of KHSRP determines its ability to slow axon growth and decrease axonal levels of mRNAs encoding growth-associated proteins.

## DISCUSSION

RBPs play critical roles in determining localized neuronal protein levels through post-transcriptional control mechanisms. Since one mRNA can generate multiple copies of protein, RNA stability needs to be tightly regulated. HuD and KHSRP both bind to ARE-containing mRNAs, with HuD stabilizing and KHSRP destabilizing the bound mRNAs. We show that PNS axons contain many different RBPs whose levels change after injury, including KHSRP. Axonal KHSRP protein levels rapidly increase in peripheral nerves following axotomy through translation of axonal *Khsrp* mRNA. This increase in axonal KHSRP likely destabilizes axonal mRNAs as *Khsrp*^*-/-*^ mice show increased axonal levels of *Gap43* and *Snap25* mRNAs, two KHSRP target mRNAs, in injured sciatic nerves compared to *Khsrp*^*+/+*^ mice. Moreover, deletion of the *Khsrp* gene accelerates regeneration of severed axons both *in vitro* and *in vivo*, indicating that intra-axonal translation of *Khsrp* mRNA slows axon regeneration. Since this effects axon regeneration from a severed axon rather than initiation of axon growth from the cell body, our results indicate that KHSRP’s role in axon growth is through its localized function in the axon. Together these data point to injury-induced increase in axonal KHSRP as an axon-intrinsic determinant of regeneration capacity.

KHSRP has been implicated in RNA splicing, trafficking and degradation as well as microRNA biogenesis (Gherzi et al., 2004; Min et al., 1997; Pan et al., 2007; Trabucchi et al., 2009). Our analyses of KHSRP target mRNA levels in sciatic nerves of the *Khsrp*^*-/-*^mice point to KHSRP’s role in RNA degradation and resulting acceleration of axon growth seen in these mice. miRNAs are clearly linked to stability of mRNAs, and their precursors (pre-miRNAs) have been shown to localize into PNS axons, with PNS nerve injury triggering localized processing of some axonal pre-miRNAs into mature miRNAs (Kim et al., 2015). However, levels of previously reported KHSRP target microRNAs (Trabucchi et al., 2009) showed no changes when comparing DRG cultures from *Khsrp*^*-/-*^ vs. *Khsrp*^*+/+*^ mice (data not shown). Transfecting the *Khsrp*^*-/-*^ DRG neurons with wild type vs. KH domain mutant KHSRP expression constructs further point to KHSRP-mediated RNA degradation rather than a secondary miRNA-mediated effect for KHSRP’s role in axon growth. KHSRP has four KH domains of about 70 amino acid residues each that can bind to single strand DNA or RNA with varying degree of specificity (Gherzi et al., 2004). The third and fourth KH domains (KH3 and KH4) are implicated in RNA decay by binding to ARE-containing mRNAs and recruiting components of the cytoplasmic exosome complex used for RNA degradation (Gherzi et al., 2004). We find both the axon growth promotion and increase in axonal *Gap43* mRNA in adult DRG neurons require an intact KH4 domain that has been shown to bind to ARE-containing mRNAs, arguing for a direct effect of KHSRP on stability of the axonal mRNAs rather than indirectly through microRNAs or other mediators. Our recent RNA-seq analyses of brain from *Khsrp*^-/-^ vs. *Khsrp*^*+/+*^ mice and mRNAs co-immunoprecipitating with KHSRP from wild type brain uncovered 527 KHSRP target mRNAs (Olguin et al., 2020). Thus, KHSRP likely binds to many other axonal mRNAs beyond *Gap43* and *Snap25* to regulate their stability. However, we cannot completely exclude indirect effects from KHSRP deletion used here, and consistent with this possibility, the work from Olguin et al. (2020) showed evidence for this with ∼1400 mRNAs that decreased in brains of *Khsrp*^*-/-*^ but showing no binding to KHSRP in RIP assays.

Several lines of evidence indicate that mRNAs can compete for interactions with RBPs and different RBPs can compete with one another for interactions with mRNAs. KHSRP as well as *Gap43* mRNA have been implicated in these competitions. We have previously shown that localized translation of *Gap43* mRNA contributes to elongating growth of DRG axons (Donnelly et al., 2013). ZBP1, in complex with HuD, is needed for axonal localization of *Gap43* mRNA (Yoo et al., 2013), and ZBP1 is also needed for axonal localization of *Actb* mRNA (Zhang et al., 2001). Interestingly, ZBP1 is expressed at limiting levels in adult DRG neurons such that *Actb* and *Gap43* mRNAs compete for localization into axons (Donnelly et al., 2013; Donnelly et al., 2011), with the axotomy-induced transcriptional increase in *Gap43* mRNA altering the stoichiometry of neuronal *Actb* to *Gap43* mRNA, and decreasing axonal localization of *Actb* mRNA (Yoo et al., 2013). *Nrn1* and *Gap43* mRNAs compete for binding to HuD with *Nrn1* mRNA’s ARE showing a higher binding affinity for HuD than *Gap43* mRNA’s ARE (Gomes et al., 2017). Furthermore, HuD and KHSRP compete for binding to *Gap43* mRNA, but they bind to the mRNA with relatively the same affinity (Bird et al., 2013). Consequently, the injury-induced increase in axonal KHSRP levels that we see here would effectively alter the localized stoichiometry of HuD to KHSRP, thereby favoring KHSRP binding to axonal *Gap43* mRNA and its subsequent degradation. Consistent with this prediction, previous RAMS analyses for proteins in crushed sciatic nerve axoplasm binding to *Gap43* mRNA’s ARE (nt 1211-1250) detected KHSRP and related KH-domain containing FUBP1 and FUBP3, but not HuD (Lee et al., 2018).

It is intriguing to speculate that the increase in *Snap25* mRNA, which was also elevated in the axoplasm of *Khsrp*^*-/-*^ mice, is similarly affected by this competition. *Snap25* mRNA contains two AREs in the 3’ UTR at nt 1085-1102 and 1586-1598 (Genbank # NM_001355254.1) that could bind to both KHSRP and HuD (Olguin et al., 2020). SNAP25 is a cytoplasmic SNARE protein involved in neurotransmitter release (Sollner et al., 1993), but SNAP25 also has reported roles in both chick retina and cortical axon development (Osen-Sand et al., 1993). Axonal translation of *Snap25* mRNA has been linked to synapse development in cortical neurons (Batista et al., 2017), so elevated axonal *Snap25* mRNA in combination with *Gap43* mRNA could contribute to the accelerated NMJ occupancy shown here in the *Khsrp*^*-/*-^ compared to *Khsrp*^*+/+*^ mice after conditioning crush lesions. With both *Gap43* and *Snap25* mRNAs linked to positive outcomes for axon growth and regeneration, these data raise the possibility that the increase in axonally synthesized KHSRP after axonal injury likely slows axon growth by decreasing the survival of mRNAs that constitute an RNA regulon supporting axon growth.

The initial increase in axonal KHSRP protein through translational upregulation of its axonal mRNA shows similar kinetics to translational induction of injury-associated mRNAs following axotomy. mRNAs encoding Importin *β*1 (*Kpnb1* mRNA), Vimentin (*Vim* mRNA), RanBP1, Stat3, mTor and Calreticulin (*Calr* mRNA) are translated in axons within the first six hours following PNS nerve crush injury (Ben-Yaakov et al., 2012; Hanz et al., 2003; Perlson et al., 2005; Terenzio et al., 2018; Yudin et al., 2008). Some of those locally synthesized injury-associated proteins help to signal retrogradely to the soma for changing gene expression, including transport of transcription factors, to support the injured neuron’s survival and axon regeneration (Rishal and Fainzilber, 2014). A retrograde calcium wave has also been shown to support axon regeneration by altering neuronal gene expression after PNS injury through epigenetic mechanisms (Cho et al., 2013). Together, these changes in gene expression likely contribute to the enhanced regeneration seen in injury-conditioned neurons. Our results showing that localized translation of *Khsrp* mRNA slows axon regeneration in injury-conditioned neurons, emphasize two critical points for the mechanisms underlying the injury-conditioning effect. First, there clearly is a response to injury conditioning that is limited to the axon since *Khsrp*^*-/-*^ injury-conditioned neurons only show enhanced axon regeneration when growth is initiated from an existing axon rather than new axon growth from the cell body. Second, this axon-intrinsic injury-conditioning effect slows axon growth, so it locally offsets the soma response to nerve injury by lowering levels of axonal mRNAs encoding growth-associated proteins. Taken together, our observations indicate that the axon growth rate seen from wild type injury-conditioned neurons is not the maximum attainable growth. However, slowing axon regeneration may be beneficial at some stages of recovery from PNS injury such as when axons reach target tissues. It is intriguing to speculate that the increase in axonal KHSRP levels as the adult animals used here begin to recover lower hind limb function after sciatic nerve crush injury may reflect such a need for axonal KHSRP.

Notably, the targeted proteomics approach used here uncovered 84 RBPs in sciatic nerve axoplasm, with many beyond KHSRP showing increased or decreased levels after nerve injury and during regeneration. Since the axoplasm isolates used here rely on extrusion in detergent-free conditions, we likely miss many RBPs that are associated with cytoskeleton or axoplasmic membrane. HuD/ELAVL4 has previously been reported to fractionate with the cytoskeleton in rat hippocampus (Pascale et al., 2004). So this may explain the decrease in HuD/ELAVL4 detected in the sciatic nerve axoplasm after injury by MS shown in Figure 1 above compared to our previous immunofluorescent analyses shown that axonal HuD/ELAVL4 levels increase during regeneration (Yoo et al., 2013). Nonetheless, considering the limitations in starting materials and sensitivity for protein detection compared to RNAs that can be amplified, we substantially increase the number of known axonal RBPs with the PRM approach. Moreover, we have uncovered a localized role for KHSRP in slowing PNS axon regeneration that is precipitated by localized translation of *Khsrp* mRNA in axons after injury. Our results expand the known functions for axonal RBPs and intra-axonal mRNA translation, and emphasize the potential to increase axon regeneration rates beyond the 1-2 mm/d typically seen after PNS injury.

## MATERIALS AND METHODS

### Animal use and survival surgery

the Institutional Animal Care and Use Committee of the University of South Carolina approved all animal procedures. Adult male Sprague Dawley rats (175-250 g) or both male and female *Khsrp* knockout (*Khsrp*^*-/-*^) (Lin et al., 2011) and wild type (*Khsrp*^*+/+*^) mice on C57/Bl6 background were used for experiments. Wild type animals were typically littermates, and heterozygous animals (*Khsrp*^*+/-*^) were used in several experiments as indicated. Isoflurane inhalation was used for anesthesia in all survival surgery experiments (see below). Animals were euthanized by CO_2_ asphyxiation at 3-7 d post-injury as indicated in results.

For peripheral nerve injury, anesthetized male rats or mice were subjected to sciatic nerve crush at mid-thigh level as previously described (Twiss et al., 2000a). For ‘double crush’ injury experiments, a unilateral peripheral sciatic nerve crush was then performed at mid-thigh level on day 0, as a ‘conditioning lesion’, and a second crush injury was performed at 0.5 cm proximal to the first crush site.

Sciatic nerve ligations were performed in male rats as described previously as the larger size of these animals provided greater precision in ligation and subsequent crush injuries (Cavalli et al., 2005). Briefly, rat sciatic nerve was ligated approximately 1 cm proximal to planned mid-thigh nerve crush site. Immediately after applying 4·0 suture, the sciatic nerve was crushed distal to the ligation site as above and then euthanized 3-16 hours later.

### Cell Culture

Dissociated cultures of adult DRGs were prepared as described (Twiss et al., 2000a). For experiments with naïve DRG neurons, all lumbar, thoracic, and lower cervical DRGs were collected. To study effects of an *in vivo* injury conditioning, L4-6 DRGs were used from ipsi-lateral (injury-conditioned) or contra-lateral (naïve) to the crush injury. DRGs were harvested in Hybernate-A medium (BrainBits, Springfield, IL) and then dissociated with collagenase as described. After centrifugation and three washes in DMEM/F12 (Life Technologies, Grand Island, NY), dissociated ganglia were cultured in complete medium containing DMEM/F12, 1 x N1 supplement (Sigma, St. Louis, MO), 10 % fetal bovine serum (Hyclone, Logan, UT), and 10 μM cytosine arabinoside (Sigma) on poly-L-lysine (Sigma) plus laminin (Millipore, Burlington, MA) coated substrates. For imaging, dissociated DRGs were cultured on coated glass coverslips. For analyses of axonal RNA levels or in vitro regeneration assay (see below), dissociated ganglia were cultured polyethylene-tetrathalate (PET) membrane inserts (1 μm pores; Falcon-Corning, Tewksbury, MA) (Zheng et al., 2001). Axons and CB were isolated from DRGs cultured on PET membranes as described (Willis et al., 2007).

For transfection, dissociated ganglia were pelleted at 100 x g for 5 min and resuspended in 100 µl ‘Nucleofector solution’ (Rat Neuron Nucleofector kit; Lonza, Alpharetta, GA). 4-6 μg of plasmid was electroporated using the AMAXA Nucleofector device (Neurons Rat DRG, G-013 program; Lonza) before plating and maintained for 48 h.

### Plasmid constructs

pAc-GFP-KHSRP and pAc-GFP-KHSRPΔKH4 constructs have been published (Bird et al., 2013). All fluorescent reporter constructs for analyses of RNA translation were based on eGFP with myristoylation element (GFP^MYR^; originally provided by Dr. Erin Schuman, Max-Plank Inst., Frankfurt) (Aakalu et al., 2001). cDNAs for the 5’UTR and 3’UTRs of *Khsrp* mRNA were custom synthesized by Integrated DNA Technologies ([IDT] Coralville, IA) and GenScript Biotech (Piscataway, NJ), respectively. The 5’UTR was engineered with 5’ *Nhe1* and 3’ *BamH1* restriction sites and cloned into pGFP^MYR^5’camk2a/3’actg (Willis et al., 2007), replacing the 5’UTR of CamK2a [GFP^MYR^5’khsrp/3’actg]. The 3’UTR sequence was engineered with 5’ *Not1* and 3’ *Xho1* restriction sites and used to replace the *Actg* 3’UTR in pGFP^MYR^5’khsrp/3’actg plasmid [pGFP^MYR^5’/3’khsrp].

### PRM-MS quantitation of RBPs

Axoplasm from 2 cm segments of sciatic nerve immediately proximal to crush site was extruded into nuclear transport buffer (20 mM HEPES [pH 7.3], 110 mM potassium acetate, 5 mM magnesium acetate) supplemented with protease/phosphatase inhibitor cocktail (Roche) and RNasin Plus (Promega, Madison, WI). Contralateral (uninjured) sciatic nerve of comparable level and length was used for control. 3 animals were used for each time point and both naïve and injured sciatic nerve axoplasm. Preparations were cleared by centrifugation at 20,000 x g, 4°C for 30 min, supernatants were diluted in 0.5 ml of TrIzol LS reagent (Invitrogen) and protein was extracted according to the manufacturer’s protocol. Protein pellets were digested with trypsin as previously described (Urisman et al., 2017).

PRM was performed on *Q Exactive Plus Mass Spectrometer* (Thermo-Fisher) online with *nanoAcquity UPLC System* (Waters, Milford, MA). Digested peptides samples (0.5 µg) were injected onto 200 cm monolithic silica-C18 column (GL Sciences, Tokyo, Japan) and separated using a 6 hour reversed phase chromatography gradient as previously described (Urisman et al., 2017). The mass spectrometer was operated in PRM mode with the following parameters: positive polarity, R = 17,500 at 200 m/z, AGC target 1e6, maximum IT 190 ms, MSX count 1, isolation window 3.0 m/z, NCE 35 %. PRM data were analyzed in Skyline v. 3.5 (MacLean et al., 2010). Skyline PRM document has been uploaded to *PanoramaWeb Public* (Sharma et al., 2018) and can be accessed at https://panoramaweb.org/axon-rbps.url.

### RNA isolation and PCR analyses

RNA was isolated from dissociated DRG neurons or cell body/axon compartments collected from insert cultures using *RNeasy Microisolation kit* (Qiagen). Sciatic nerve was cut in small pieces and digested with collagenase at 37°C for 30 min with intermittent trituration. RNA was isolated from the collagenase-treated nerve using Trizol LS reagent (Invitrogen, Carlsbad CA) according to manufacturer’s instructions. RNA concentration was measured by fluorimetry with *Ribogreen* (Life Technologies) and 10-50 ng of RNA was reverse transcribed with *Sensifast cDNA synthesis kit* (Bioline, London, UK). DRG axonal purity was assessed by RT-PCR, performed with primers designed to detect cell body-restricted mRNAs (*cJun* and *Map2*) and glial cell-specific mRNAs (*Gfap*). ddPCR was performed according to manufacturer’s procedure with *Evagreen* detection (Biorad, Hercules, CA). Mitochondrial 12S ribosomal RNA (*Mtrnr1*) and Hmgb1 mRNA levels were used for normalizing RNA yields across different isolates. Following primers were used RT-PCR and RTddPCR (all from IDT, listed as 5’ to 3’): *Mtrnr1*, sense – GGCTACACCTTGACCTAACG and antisense – CCTTACCCCTTCTCGCTAATTC; *Actb*, sense – CTGTCCCTGTATGCCTCTG and antisense – ATGTCACGCACGATTTCC; *cJun*, sense – GCAAAGATGGAAACGACCTTCTAC and antisense – AAGCGTGTTCTGGCTATGC *Gfap*, sense – AGTTACCAGGAGGCACTTG and antisense – GGTGATGCGGTTTTCTTCG; *Hmgb1*, sense – CATGGGCAAAGGAGATCC and antisense – CTCTGAGCACTTCTTGGAG; *Gap43*, sense – CAGGAAAGATCCCAAGTCCA and antisense – GAGGAAAGTGGACTCCCACA; *Map2*, sense – CTGGACATCAGCCTCACTCA and antisense – AATAGGTGCCCTGTGACCTG; and, *Snap25*, sense – CAAATTTAACCACTTCCCAGCA and antisense – CAGAATCGCCAGATCGACAG.

### Immunofluorescent staining

Immunofluorescence was performed as previously described with all steps at room temperatures unless specified otherwise. Coverslips were fixed with 4% paraformaldehyde (PFA) in phosphate-buffered saline (PBS) for 15 min at room temperature and washed 3 times in PBS. PBS washed neurons were permeabilized with 0.3 % Triton X-100 in PBS for 15 min and then blocked in 5 % BSA for 1 h. Neurons were incubated with primary antibodies overnight in humidified chambers at 4°C. Primary antibodies consisted of: chicken anti-NFH, NFM and NFL cocktail (1: 500; Aves Lab, NFH # AB_2313552, NFM # AB_2313554, and NFL # AB_2313553), RT97 mouse anti-NF (1:500; Devel. Studies Hybridoma Bank, Iowa City, IA). After washes in PBST, coverslips were incubated with combination of FITC-conjugated donkey anti-mouse, Cy5 conjugated donkey anti-chicken (both at 1:500; Jackson ImmunoRes., West Grove, PA) as secondary antibodies for 1 h. After 1 h, coverslips were washed 3 times in PBS, rinsed with distilled H_2_O, and mounted with *Prolong Gold Antifade with DAPI* (Life Technologies).

For regeneration studies on mouse sciatic nerve and quantifying axonal content of KHSRP *in vivo*, sciatic nerve segments were fixed for 4 h in 4 % PFA and then cryoprotected overnight in 30 % sucrose in PBS at 4°C. 10 µm cryostat sections for rat sciatic nerve and 20 µm cryostat sections for mouse sciatic nerve were processed for immunostaining as previously described (Merianda et al., 2013). Primary antibodies consisted of RT97 mouse anti-NF (1:500), rabbit anti-KHSRP (1:200; Novus Biologicals, NBP1-8910, Centennial CO), rabbit anti-Stathmin-2/SCG10 (1:500; Novus Biologicals, NBP1-49461). Stathmin-2/SCG10 immunofluorescence was used to detect regenerating mouse sciatic nerve axons (Shin et al., 2012). Cy3-conjugated donkey anti-rabbit and FITC-conjugated donkey anti-mouse in combination were used as secondary antibodies for rat sciatic nerve (both at 1:500, Jackson ImmunoRes). Cy3-conjugated donkey anti-rabbit antibodies were used on mice sciatic nerve (1:500, Jackson ImmunoRes).

All samples were mounted with *Prolong Gold Antifade* and imaging was performed at room temperature. Samples were analyzed by either epifluorescent or confocal microscopy. Leica DMI6000 epifluorescent microscope with ORCA Flash ER CCD camera (Hamamatsu) was used for epifluorescent imaging. Confocal imaging for immunofluorescence was performed on a Leica SP8X microscope (DMI6000 M platform; Buffalo Grove, IL) fitted with a galvanometer Z stage and HyD detectors; HC PL Apo 63x/1.4 NA objective (oil immersion) was used with acquisition parameters matched for individual experiments using LAS-X software. Z-stack images were post-processed by Leica *Lightning Deconvolution* integrated into *LASX* software. Deconvolved image stacks were projected into single plane images using the maximum pixel intensities.

### Neuromuscular Junction staining

NMJ staining protocol was performed as previously published with minor modifications (Maimon et al., 2018). Briefly, all steps were carried out at room temperature. Gastrocnemius muscle (GM) was cleared of any connective tissue, washed in PBS, fixed in 4 % PFA, washed in PBS 3 times for 5 min each. Muscle was then dissected into smaller pieces and incubated with 1 μg/ml of Alexa Fluor 488 conjugated α-Bungarotoxin for 4 h with rocking (Thermo-Fisher, B13422). Tissues were washed with PBS 3 times for 5 min each, treated with methanol at −20°C for 5 min, and rinsed in PBS 3 times for 5 min each with rocking. Tissues were blocked for 1 h with 2 % BSA, 0.4 % Triton X-100 in PBS. Tissues were then incubated overnight with the following cocktail of primary antibodies to presynaptic components diluted in blocking solution with rocking: rabbit anti-NF 200 (1:200; Sigma, N4142), mouse anti-synaptophysin (1:300; Millipore, MAB5258), rabbit anti-synapsin-I, (1:200; Millipore, AB1543P). The following day, tissues were rinsed 3 times 5 min each in PBS while rocking. Samples were then incubated in Cy3-conjugated donkey anti-rabbit and Cy3-conjugated donkey anti-mouse antibodies (1:500; Jackson ImmunoRes) for 4 h. After rinsing in PBS, muscle fibers were spread into monolayers under a stereomicroscope and affixed to slides using *Prolong Gold Antifade*; coverslips were sealed with clear nail polish.

Leica SP8X microscope as above was used for imaging. Sequential scanning was used to separate the green and red channels and Z stack with 200 nm interval using a HC PL Apo 63x/1.4 NA oil immersion objective.

### Fluorescence in-situ hybridization (FISH)

smFISH plus IF was used to detect *Khsrp* mRNA in DRG and sciatic nerve. We used custom designed Cy3-labelled Stellaris probes (LGC Biosearch Tech, Middlesex, UK) for mouse *Khsrp* mRNA (Genbank ID # NM_010613.3) with Cy3-labelled scramble probe for control. RT97 mouse anti-NF (1:200) was used as a primary antibody and FITC-conjugated donkey anti-mouse (1:200) was used as secondary. Samples processed without addition of primary antibody served as control for antibody specificity. Samples were mounted with *Prolong anti-fade*.

smFISH/IF on DRG cultures was performed as described previously (Kalinski et al., 2015). Briefly, coverslips were rinsed in PBS and then fixed in buffered 2 % paraformaldehyde for 15 min, with all steps carried out at room temperature unless specified otherwise. Coverslips were rinsed 2 times in PBS, then permeabilized in 0.3 % Triton X-100 in PBS for 5 min. Samples were equilibrated for 5 min in hybridization buffer (50 % dextran sulphate, 10 μg/ml *E. coli* tRNA, 10 mM ribonucleoside vanadyl complex, 80 µg BSA, and 10 % formamide in 2X SSC), and then incubated with 12.5 µM probe plus mouse anti-NF (1:200) for 12 h at 37°C. Coverslips were then washed in PBS + 0.3 % Triton X-100 3 times, followed by incubation with FITC-conjugated donkey anti-mouse for 1 h. After rinse in PBS, post fixation in 2 % PFA for 15 min, and a second PBS wash, coverslips were inverted and mounted on glass slides.

Detection of mRNA in mouse tissues was done as previously described (Kalinski et al., 2015), with all steps carried out at room temperature unless specified otherwise. Briefly, sciatic nerve segments were fixed for 4 h in 2 % PFA, cryoprotected overnight in 30 % buffered sucrose at 4°C, and then cryosectioned at 20 µm thickness (sections were stored at −20°C until used). Sections were brought to room temperature, washed three times in PBS for 5 min each, and then treated with 20 mM glycine and fresh 0.25 M NaBH_4_ in PBS (3 times, 10 min each for both) to quench autofluorescence. Sections were quickly rinsed in 0.1 M Triethylamine (TEA) and then incubated in 0.1 M TEA + 0.25 % Acetic Anhydride for 10 min. Sections were dehydrated in 70, 95, and 100 % ethanol (3 min each) and then delipidated in chloroform for 5 min followed by 100 and 95 % ethanol (3 min each). After washing in 2X SSC, sections were incubated overnight at 37°C in a humidified chamber with 12.5 µM probe and RT97 anti-NF (1:100) in hybridization buffer. The following day, sections were washed in 2X SSC + 10 % formamide at 37°C for 30 min, followed by two incubations in 2X SSC for 5 min each. Sections were then briefly rinsed in PBS + 1 % Triton-X100, and then incubated for in donkey anti-mouse FITC antibody diluted in 10 X blocking buffer (1:100; Roche) + 0.3 % Triton-X100 for 1 h. Sections were finally washed in PBS for 5 min, post-fixed in 2 % PFA for 15 min, washed 3 times in PBS (5 min each), rinsed in DEPC-treated water, and mounted under glass coverslips. smFISH and IF signals were imaged using Leica SP8X as above. 63X/NA 1.4 objective and pulsed white light laser was used for imaging RNA in both culture and tissue samples. Scramble probe was used to set the image acquisition parameter that would not acquire any nonspecific signal from scramble probe. Taking XYZ image stacks at least two locations in each section scanned nerve sections.

### Detection of nascently synthesized proteins

Rat sciatic nerve from three animals per condition was either left naïve or *in vitro* crushed and incubated in DMEM medium containing, 10 % FBS + Cyclosporin A (20 µM; Sigma) + 1 % penicillin/streptomycin. Nerves were treated with 200 µg/ml anisomycin or vehicle (0.1 % DMSO) for 3 h at 37°C, followed by adding 100 µg/ml O-propargyl-puromycin (OPP; Invitrogen) for 1 h at 37°C. Axoplasm was extracted in 1 mL of transport buffer (20 mM HEPES [pH 7.4], 100 mM sodium acetate, 5 mM magnesium acetate), after extraction SDS added to 1 %. Protein concentration was quantified by BCA assay and 350 µg of total protein was used for biotin conjugation by click chemistry (100 µM biotin-PEG3-azide) according to manufacturer’s instruction. The reaction mix was incubated for 2 h at room temperature on a rotator. 5 volumes ice-cold acetone was added to precipitate the protein. Protein pellets were resuspended in PBS containing 1 % SDS. Streptavidin pull-down was carried out overnight at 4°C in 1 ml volume containing 60 μl of streptavidin magnetic beads (Life Technologies), 1 % NP40, 0.1 % SDS and 1x *Complete EDTA-free protease inhibitor cocktail* (Roche) in PBS. 10 % of protein used for pull-down was taken to generate input samples. Beads were washed three times for 10 min with 1 % NP40, 0.1 % SDS in PBS at room temperature. Proteins were eluted from streptavidin beads by boiling for 10 min in 2x Laemmli sample buffer and then adjusted to 1x with PBS for denaturing polyacrylamide gel electrophoresis (SDS/PAGE).

### Immunoblotting

Adult mouse DRG cultures (3 days *in vitro*, ∼80,000 neurons/well) were lysed in Laemmli sample buffer and denatured by boiling at 95°C x 5 min. Rat axoplasm from naïve and crushed sciatic nerves was extruded in nuclear transport buffer (20 mM HEPES [pH 7.3], 110 mM potassium acetate, 5 mM magnesium acetate, supplemented with protease inhibitors) as previously described (Hanz et al., 2003). Lysates were cleared of debris by centrifugation at 15,000 x g for 15 min at 4°C and then normalized for protein content using Bradford assay (BioRad). Normalized protein lysates were fractionated by 10 % SDS/PAGE and transferred onto a PVDF membrane (GE Healthcare Life Sciences, Marlborough, MA). After blocking in 5% non-fat dried milk powder (Biorad) diluted in Tris-buffered saline with 1% Tween 20 (TBST), membranes were probed overnight at 4°C with Rabbit anti-KHSRP (1:1000; Novus) or Rabbit anti-GAPDH (1:2000; Cell Signaling Tech, Beverly, MA), Streptavidin-HRP (1:10000, Abcam) antibodies diluted in blocking buffer. Blots were washed in TBST and then incubated with horseradish peroxidase (HRP)-conjugated anti-rabbit IgG (1:2000; Cell Signaling Tech) diluted in blocking buffer for 1 h at room temperature. Blots were washed in TBST and signals were detected by ECL Prime^™^ (GE Healthcare Life Sciences).

### Fluorescence recovery after photobleaching

FRAP experiments were conducted at 37°C, 5 % CO_2_ as previously described (Yudin et al., 2008). Briefly, dissociated adult mouse DRG cultures transfected with the GFP^MYR^5’/3’khsrp were equilibrated in culture medium as above except phenol red was excluded. A region of interest (ROI) in the most distal axon of dissociated DRG neurons was photobleached with 488 nm argon laser set at 100 % power for 80 frames at 0.65 sec each. Pre-bleach and post-bleach signals were captured using 70 % power for 488 nm laser line every 30 sec (2 for pre-bleach and 30 for post-bleach). Leica SP8X confocal microscope was used for imaging at 37°C, 5% CO_2_ with 63X/NA 1.4 objective. Pinhole was set to 3 Airy units for pre-bleach, bleach, and post-bleach sequences to ensure full thickness excitation of the axon. ROIs were 40 x 40 µm and at least 250 µm from the soma.

### Image analyses

*Image J* was used to quantify protein and RNA levels in sciatic nerve tissues from optical planes of XYZ scans. Axon only signal was extracted via *Colocalization plug-in* that extracted only protein or RNA signals that overlapped with axonal marker (NF) in each plane. Extracted ‘axon only signal ‘was projected as a separate channel (Kalinski et al., 2015). Signal intensities were then calculated from each XY plane of these axon only channels. NF immunoreactivity area was used to normalize signal intensities across the individual XY planes. The relative signal intensity was then averaged for all tiles in each biological replicate.

To assess regeneration *in vivo*, tile scans were post-processed by *Straighten plug-in* for *ImageJ* (http://imagej.nih.gov/ij/). SCG10 fluorescence intensity was measured along the length of the nerve using *ImageJ*. Regeneration index was calculated by measuring the average SCG10 intensity in bins across at least 3 mm distal to the crush site. The crush site was defined by the position along the nerve length with maximal SCG10 intensity (secondarily confirmed by DAPI signals and DIC images).

For analyses of axon growth *in vitro*, dissociated DRGs were immunostained with NF antibodies as described above. Images from 36 or 48 h cultures were used for neurite length and branching parameters (neurites/cell body and branch density) using WIS-Neuromath (Rishal and Fainzilber, 2014).

For NMJs, confocal Z stacks from muscle were projected as single XY images using *ImageJ*. Nerve terminal and endplate (AchRs) areas were calculated by *Image J* and fractional occupancy of NMJ was calculated by dividing nerve terminal area to endplate AChR area as described (Maimon et al., 2018).

For FRAP image analyses, raw images sequences were analyzed for recovery in the bleached ROI using Leica confocal software package. Recovery was determined relative to pre-bleach and post-bleach signals, which were set at 100 and 0 % to allow for comparisons between experiments and between neurons.

### Statistical analyses

*GraphPad Prism* software package (La Jolla, CA) was used for statistical analyses. One-way ANOVA with Tukey post-hoc was used to compare between data points and Student’s t-test was used to compare two independent groups for most experiments. For FRAP studies, two-way ANOVA with Tukey post-hoc was used, where control values were compared to cycloheximide- and anisomycin-treated cultures for each time point. All experiments were performed in at least triplicate. P ≤ 0.05 was considered as statistically significant..

## Supporting information

Supplemental Methods - Reagent Table

## Acknowledgements

This work was supported by grant awards from Wings for Life Spinal Cord Injury Research Foundation (WFL-US-09/18 to PP), National Institutes of Health (R01-NS089633 to JLT and NPB; K01-NS105879 to TPS), the Dr. Miriam and Sheldon G. Adelson Medical Research Foundation (to JLT and ALB), South Carolina Spinal Cord Injury Research Fund (2019 PD-02 to PKS), and SC EPSCoR Stimulus Grant Program (18-SR04 to JLT). JLT is the incipient South Carolina SmartState Chair in Childhood Neurotherapeutics at the University of South Carolina.

## Conflict of Interest

The authors declare no financial conflicts.

## Notes

https://panoramaweb.org/axon-rbps.url

### Competing Interest Statement

The authors have declared no competing interest.

