## Supplemental Figures for "Intra-axonal translation of *Khsrp* mRNA slows axon regeneration by destabilizing localized mRNAs"

**SUPPLEMENTAL FIGURE LEGENDS**

**Supplemental Figure S1**: ***PRM analyses of PNS axoplasm RNA binding proteins.*** Median $Log_2$ fold-change for each protein from Figure 1A across the different time points post-injury relative to naïve axoplasm is shown. Red lines indicate statistically significant differences across the time points.

**Supplemental Figure S2**: ***Loss of KHSRP has no effect on axon branching.***

Number of axons extending from each soma (**A**) and branch density along axons (**B**) are shown for naïve and 7 d injury-conditioned L4-6 DRGs cultures from *Khsrp*$^{+/+}$ and *Khsrp*$^{-/-}$ mice as mean ± SEM (≥ 75 neurons analyzed per sample over 3 independent cultures; no significant differences detected by one-way ANOVA with Tukey's multiple comparison test).

**Supplemental Figure S3**: ***Deletion of KHSRP modestly increases axon regeneration at 7 days after crush injury.***

Regeneration indices calculated as in Figure 3B are shown for *Khsrp*$^{+/+}$ and *Khsrp*$^{-/-}$ mice sciatic nerves at 7 days after a single nerve crush injury (N = 5 mice per genotype and condition; no significant differences by two-way ANOVA with Bonferroni post-hoc analysis).

**Supplemental Figure S4**: ***Nerve-intrinsic elevation of axonal KHSRP distal to a ligation is predominantly at the crush site.***

**A)** Representative immunofluorescent images for ligated nerves as in Figure 5A (proximal = left; distal = right) show accumulation of APP proximal to the ligation and Stat3 distal to the ligation [Scale bar = 100 µm].

**B-C)** Representative confocal image pairs for KHSRP (grey) + NF (magenta) and axonal KHSRP (spectral intensity) as in Figure 5B are shown for naïve (**B**) and ligation proximal + ligation distal (**C**, proximal = left; distal = right) sciatic nerve [Scale bar = 5 µm].

__Supplemental Figure S5__: ***FRAP analyses for local translation of Khsrp 5'/3' reporter mRNA in DRG axons.***

Representative time lapse sequence for $GFP^{MYR}$5'/3 'expressing DRG neurons under control (upper row) and cycloheximide treatment (lower row). The boxes represent the bleached ROI that shows attenuated recovery when protein synthesis is inhibited [Scale bar = 10 µm].

### Supplemental Figure S1

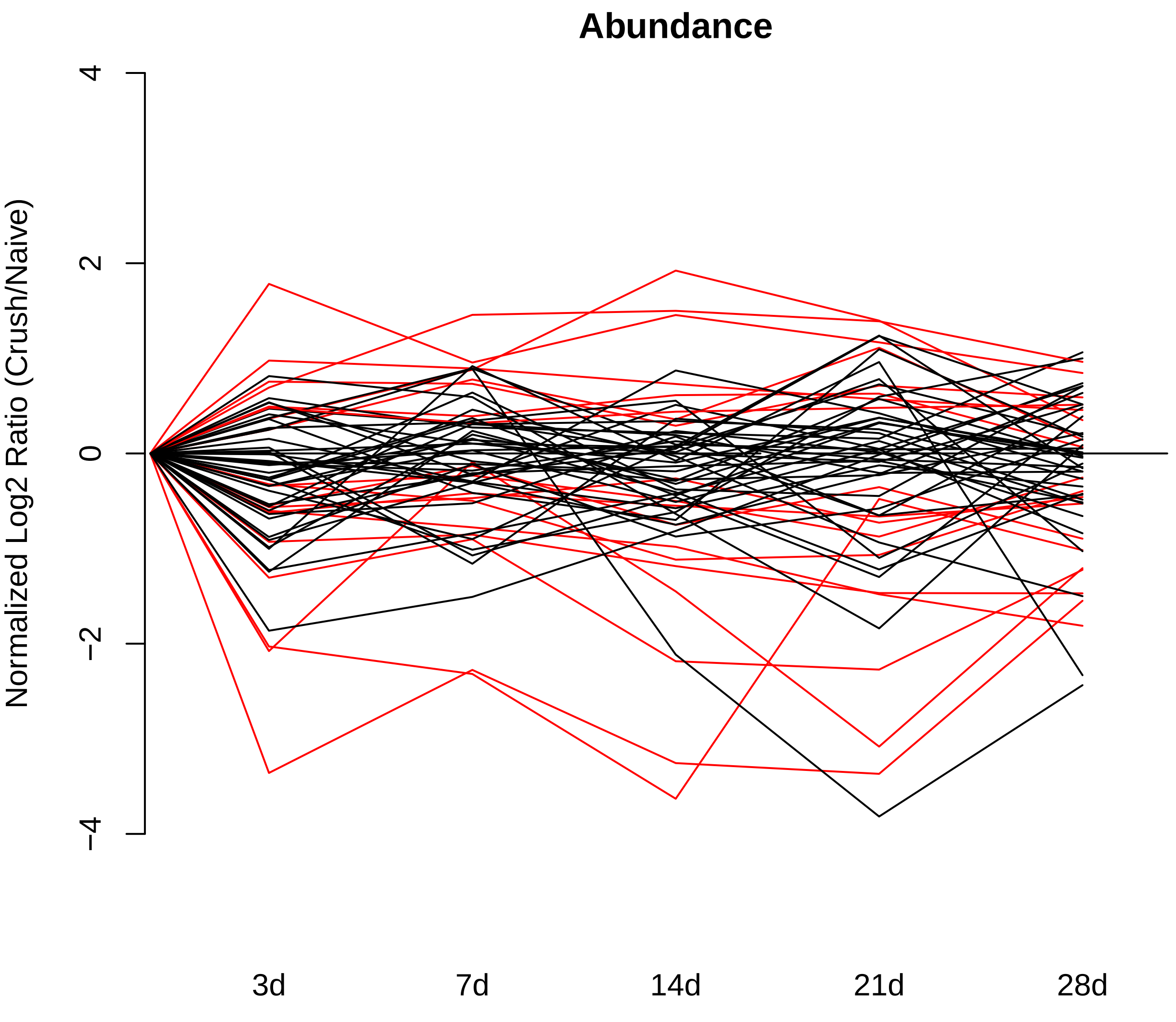

**Supplemental Figure S2**

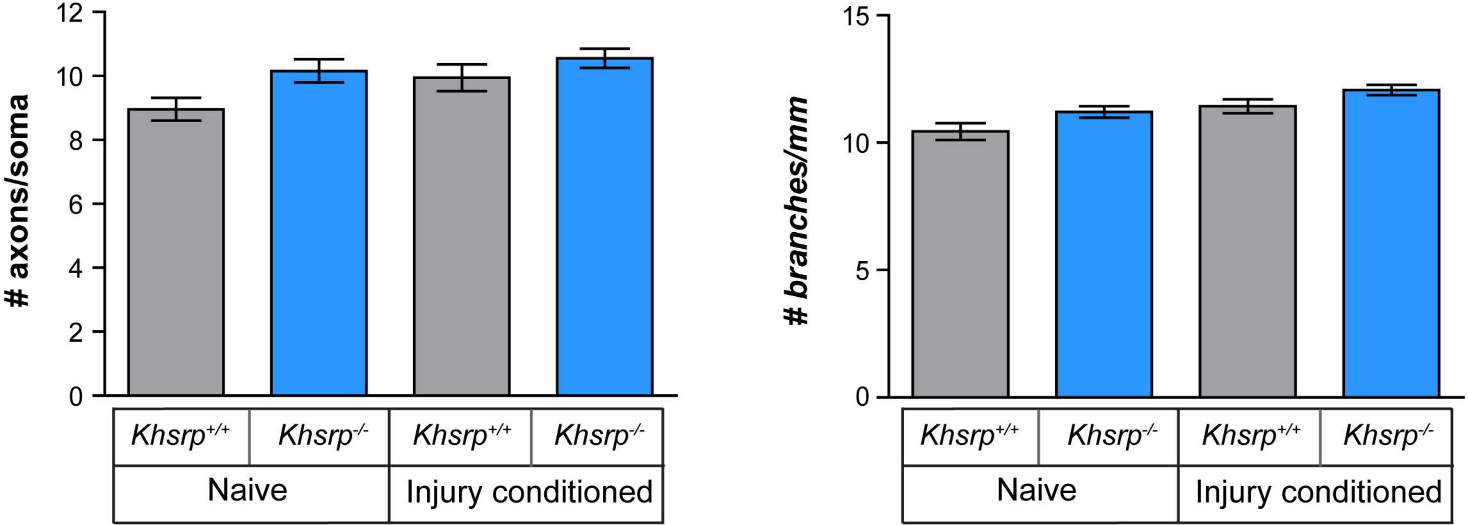

#### Supplemental Figure S3

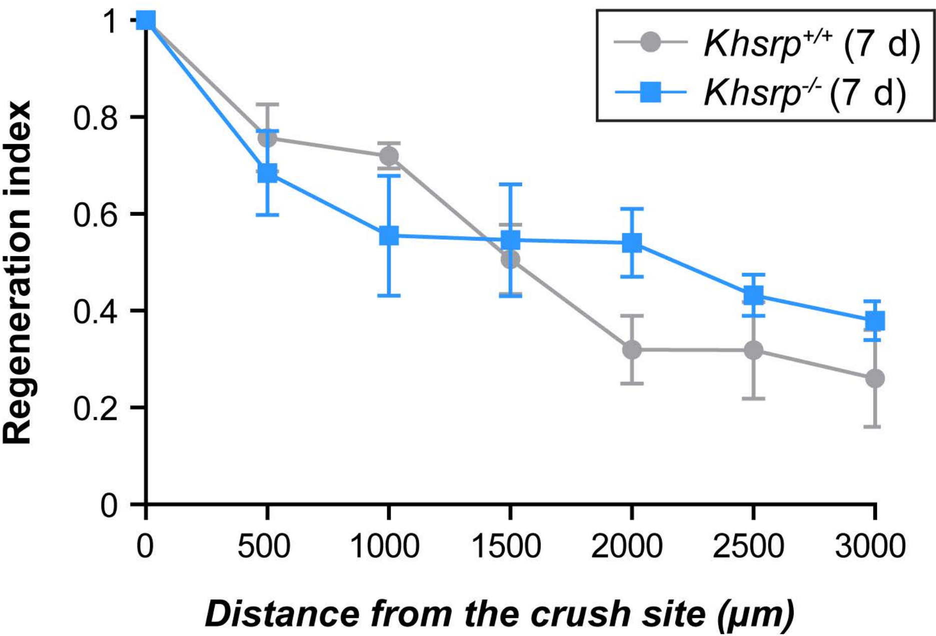

### Supplemental Figure S4

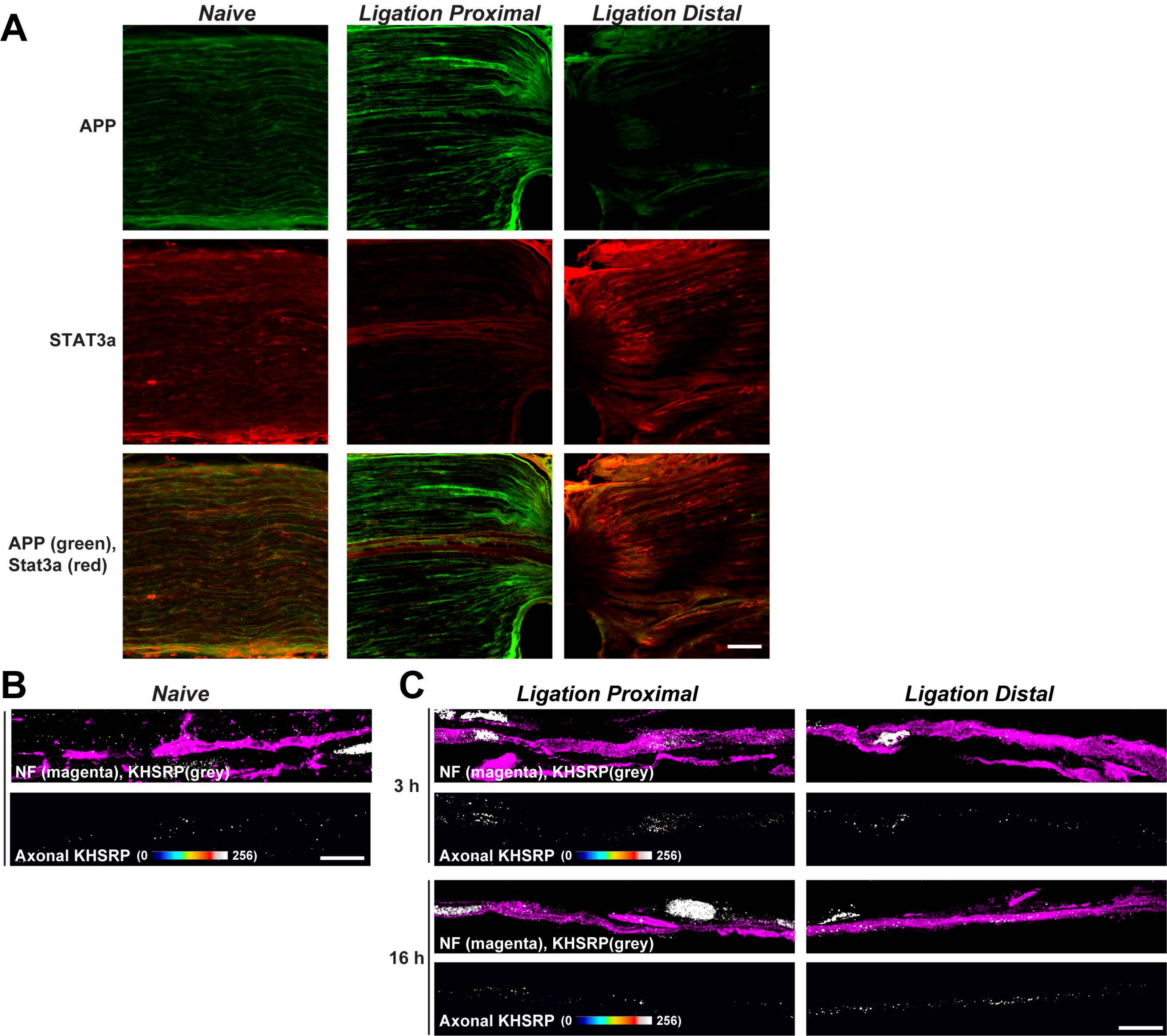

**Supplemental Figure S5**

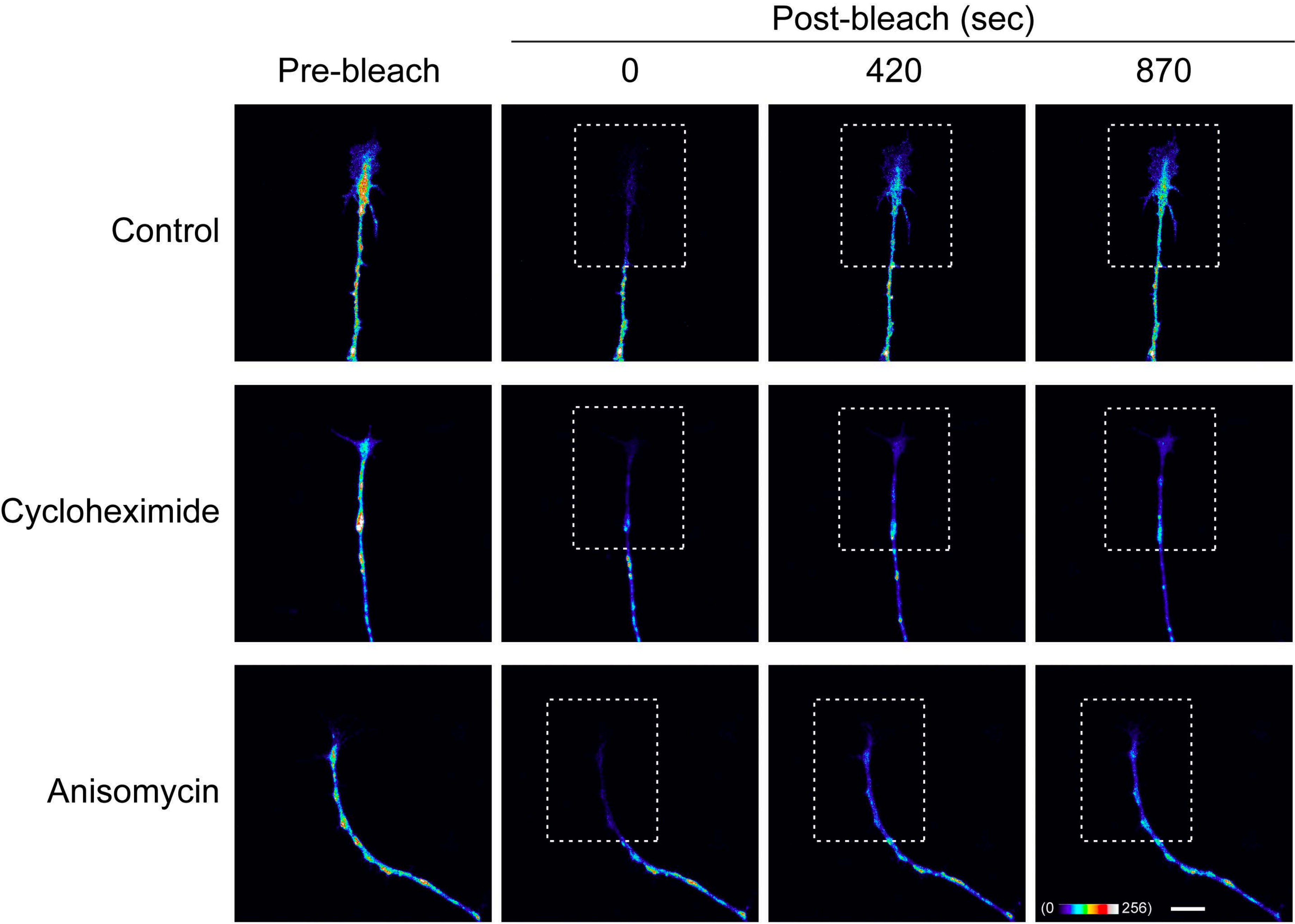
