## Supplemental Methods - Reagent Table for "Intra-axonal translation of *Khsrp* mRNA slows axon regeneration by destabilizing localized mRNAs"

### KEY RESOURCES TABLE

| REAGENT or RESOURCE | SOURCE | IDENTIFIER |
|---|---|---|
| **Antibodies** | | |
| Rabbit anti-KHSRP | Novus Biologicals | Cat # NBP1-8910 |
| Rabbit anti-Stathmin-2/SCG10 | Novus Biologicals | Cat # NBP1-49461 |
| rabbit anti-GAPDH | Cell Signaling Technology | Cat # 2118S |
| RT97 mouse anti-neurofilament | Devel. Studies Hybridoma Bank | Cat # AB_528399 |
| Chicken anti-Neurofilament (heavy) | Aves | Cat # NFH |
| Chicken anti-Neurofilament (medium) | Aves | Cat # NFM |
| Chicken anti-Neurofilament (light) | Aves | Cat # NFL |
| Rabbit anti-Neurofilament 200 (NF200) | Sigma | Cat # N4142 |
| Mouse anti-Synaptophysin, clone SY38 | EMD Millipore | Cat # MAB 5258 |
| Rabbit anti- Synapsin | Millipore | Cat # AB1543P |
| Cy3-conjugated donkey anti-mouse | Jackson ImmunoRes | Cat # 715-165-151 |
| FITC-conjugated donkey anti-rabbit | Jackson ImmunoRes | Cat # 711-095-152 |
| Cy3-conjugated donkey anti-rabbit | Jackson ImmunoRes | Cat # 711-165-152 |
| FITC-conjugated donkey anti-mouse | Jackson ImmunoRes | Cat # 715-095-151 |
| Cy5-conjugated donkey anti-chicken | Jackson ImmunoRes | Cat # 703-175-155 |
| Alexa Flour 488 conjugated α-bungarotoxin | Thermofisher Scientific | Cat # B13422 |
| HRP conjugated anti Rabbit IgG | Cell Signaling Technology | Cat # 7074 |
| **Bacterial and Virus Strains** | | |
| Not Applicable | | |
| Biological Samples | | |
| DRGs and Sciatic Nerve isolated from SD rats and *KHSRP*$^{+/+}$, *KHSRP*$^{+/-}$ and *KHSRP*$^{-/-}$ mice | | |
| **Chemicals, Peptides, and Recombinant Proteins** | | |
| Cyclohexamide | Sigma | Cat # C7698 |
| Anisomycin | Sigma | Cat # A5862 |
| Cyclosporin A | TCI Chemicals | Cat # C2408 |
| O-propargyl-puromycin | Thermo Fisher Scientific | Cat # C10459 |
| Biotin Azide | Thermo Fisher Scientific | Cat # B10184 |
| **Critical Commercial Assays** | | |
| Click-IT™ Biotin Protein Analysis Detection Kit | Life Technologies | Cat # C33372 |
| Trizol LS reagent | Thermo Fisher Scientific | Cat # 10296028 |
| N1 media supplement | Sigma | Cat # N6530-5ML |
| **Deposited Data** | | |
| Not applicable | | |
| **Experimental Models: Cell Lines** | | |
| Not applicable | | |
| **Experimental Models: Organisms/Strains** | | |
| Rats: Sprague Dawley strain | Sprague Dawley | |
| Mice: *Khsrp*$^{-/-}$ on C57Bl/6 background | (Lin et al., 2011) | |

| Oligonucleotides | | |
|---|---|---|
| Stellaris probes against mouse *Khsrp* | Biosearch Tech | Cat # SMF1063-5 |
| *Mtrnr1* – sense, GGCTACACCTTGACCTAACG; anti-sense, CCTTACCCCTTCTCGCTAATTC | IDT | |
| *Actb* – sense, CTGTCCCTGTATGCCTCTG; anti-sense, ATGTCACGCACGATTTCC | IDT | |
| *cJun* – sense, GCAAAGATGGAAACGACCTTCTAC; anti-sense, AAGCGTGTTCTGGCTATGC | IDT | |
| *Gfap* – sense, AGTTACCAGGAGGCACTTG; anti-sense, GGTGATGCGGTTTTCTTCG | IDT | |
| *Hmgb1* – sense, CATGGGCAAAGGAGATCC; anti-sense,CTCTGAGCACTTCTTGGAG | IDT | |
| *Gap43* – sense, CAGGAAAGATCCCAAGTCCA; anti-sense, GAGGAAAGTGGACTCCCACA | IDT | |
| *Map2* – sense, CTGGACATCAGCCTCACTCA; anti-sense, AATAGGTGCCCTGTGACCTG | IDT | |
| *Snap25* – sense, CAAATTTAACCACTTCCCAGCA; antisense,CAGAATCGCCAGATCGACAG | IDT | |
| **Recombinant DNA** | | |
| $GFP^{MYR}$5'khsrp/3'khsrp | This paper | |
| pAc-GFP-KHSRPΔKH4 | (Bird et al., 2013) | |
| pAc-GFP-KHSRP | (Bird et al., 2013) | |
| pAc-GFP | (Bird et al., 2013) | |
| **Software and Algorithms** | | |
| Graphpad Prism | https://www.graphpad.com | |
| ImageJ | https://imagej.nih.gov/ij/ | |
| ImageJ Colocalization Plugin | https://imagej.nih.gov/ij/plugins/colocalization.html | |
| Neurolucida | https://www.mbfbioscience.com/neurolucida | |
| **Other** | | |
